# Neuromuscular Architecture of the Siphonophore colony

**DOI:** 10.64898/2026.08.03.742546

**Authors:** Tigran P. Norekian, Leonid L. Moroz

## Abstract

Siphonophores are colonial hydrozoans with unprecedented differentiation and specialization, in which individual zooids are transformed into ‘functional organs’ rather than autonomous polyps capable of feeding. As a result, the entire colony acts as a single, modular–individual with the highest level of coordination and integration, from development through behavior. Deciphering these integrative mechanisms requires understanding the microanatomical organization of the nervous system in all elements of the colony. Here, using two immunohistochemical markers (anti-tubulin and anti-RFamide antibodies), we systematically characterize the neural systems across the entire *Nanomia* colony, encompassing pneumatophore, stem and all zooid classes (nectophores, gastrozooids, palpons, male and female gonophores, and protective zooids). The use of two neuronal markers enables visualization of distinct neural subpopulations, some of which are not revealed by a single marker. We provide evidence of neuroanatomical interactions within all elements of the colony, including contributions of giant axons, stem polygonal networks, and RFamide-ir neural rings at the base of each zooid, as well as describe different subpopulations of neural networks in the body of various zooids. The presented mapping facilitates identification of novel conductive and signaling pathways for future analysis of the cellular basis of behavioral integration within decentralized, broadly distributed networks and non-neuronal elements of these unique superorganisms.

## INTRODUCTION

*The achievement of siphonophores is “one of the greatest in the history of evolution” (Wilson, 1975)*.

Order Siphonophorae (class Hydrozoa, phylum Cnidaria) includes about 180 identified species (Church et al., 2025; Claver et al., 2024; Mapstone, 2014; Munro et al., 2018). Siphonophores represent “something of a paradox, being colonies in the phylogenetic sense but developing and behaving as individual organisms” (Mackie, 1978). According to Wilson (1975) “the resolution of the paradox is that siphonophores are both organisms and colonies”. In a phylogenetic sense, they are colonies composed of dozens or hundreds of zooids, reaching up to 40 meters in size, larger than a blue whale (Dunn, 2009). However, each of these colonies develops and behaves like an individual, with remarkable coordination and extraordinary division of labor among the batteries of different zooids (Dunn, 2005, Mackie et al., 1988), functioning together as a super-organism (Mackie, 1963).

G. Mackie, a pioneer in siphonophore neurobiology, stressed: “they originated as colonies, but have *evolved* as individuals,” progressing further in terms of system-level and behavioral integration than any other animal colonies (Mackie, 1978). In this evolutionary trajectory, siphonophores have achieved the most “advanced grade of construction by making organs out of what were originally individual zooids” (Mackie, 1978). Siphonophores use the same genome to transform polyps into specialized organs, as genetic twins, with sustained division of polyp’s structure and functions across the entire colony (Dunn, 2009). They succeed as superorganisms by avoiding the inherent conflict between individuality by efficiently sharing nutrients and information among all zooids, using an elongated stem as an attachment and coordination center for all individuals. Such a unique organization requires a unique system-level coordination (Dunn and Wagner, 2006) both in development and behavioral integration (Du Clos et al., 2022; Mackie et al., 1988; Strock et al., 2023; Costello et al., 2015; Sutherland et al., 2019b). However, we only begin to understand these system-level integrative mechanisms.

The genus *Nanomia* (Hosia et al., 2024) became an accessible reference preparation to obtain critical information on its zooid organization (Church et al., 2015), stem cells (Siebert et al., 2015), differential gene expression (Ahuja et al., 2024, Munro et al., 2022, Siebert et al., 2011), and deciphering the cellular bases of siphonophore behaviors (Mackie, 1978, Mackie et al., 1988, Norekian and Meech, 2020, Norekian and Meech, 2026). *Nanomia* exhibits a highly integrated avoidance behavior and directional swimming patterns (Costello et al., 2015; Sutherland et al., 2019a), in which the neural and epithelial pathways play distinct roles in signal propagation (Mackie, 1964, Mackie, 1965, Mackie, 1973, Mackie, 1978, Mackie et al., 1988, Spencer, 1971). Giant axons and ectodermal neural networks in the *Nanomia* stem were originally described using electron and phase-contrast light microscopy (Mackie, 1973, Mackie, 1978). Comparable neural structure in the stem including a pair of giant axons in the ectoderm and a superficial neural plexus have been found in a calycophore siphonophore *Chelophyes* (Mackie and Carre, 1983). Later, custom-made antiserum to the Arg-Phe-amide (RFamide) sequence was used to stain the nervous systems of *Nanomia bijuga* and other physonectid siphonophores such as *Agalma elegans*, *Forskalia edwardsi*, *Forskalia leuckarti*, and *Halistemma rubrum,* (Grimmelikhuijzen et al., 1986). That work revealed as RFamide-positive, the following structures: ectodermal nerve net in the stem and pneumatophores, scattered net of neurons in gastrozooids and tentacles, nerve rings at the base of gastrozooids, palpons, and gonozooids, transverse bands (collars) along the stem, and immunoreactive ring at the margin of nectophores (Grimmelikhuijzen et al., 1986). In *Nanomia,* similar results were obtained for the expression of a gene encoding endogenous *nb-rfamide* peptide using *in situ* hybridization (Church et al., 2015). Although *in situ* hybridization provides highly specific mapping of cell somata, the distribution of neural processes and network identification is challenging because neuropeptide mRNA does not transport to distant processes.

Recently, a morphological and functional characterization of nectophores and the nectosomal stem integration was conducted using tubulin and FMRFa immunoreactivity and electrophysiological recordings (Norekian and Meech, 2020, Norekian and Meech, 2026). Specifically, these authors described the nectophore nervous system, including the ring nerve, the sensory upper nerve, and the lower nerve, which ends in the small terminal ganglion (Norekian and Meech, 2020). The terminal ganglion comes into close contact with the cone-shaped protrusions of the stem, which serve as nectophore docking stations, and with the polygonal neural networks that cover the entire surface of those cones and the rest of the stem, and consist of both tubulin-and FMRFa-immunoreactive neural subpopulations – thus providing the morphological foundation for neuronal coordination between the individual nectophores and the stem (Norekian and Meech, 2026). Although the nectosome region and integration between individual nectophores were studied in detail, the rest of the colony remained less explored and required more focused attention. Integration and validation of different methods and approaches are also needed.

Here, we provide a detailed map of the neural and muscular organization across the entire *Nanomia* colony, representing all zooid classes. The anatomical structure of the nervous system in the nectosome region is briefly overviewed, with some new details added, while more focus is placed on the siphosome region (Fig. 1). We used two different markers to identify various subpopulations of neural elements in *Nanomia* zooids – anti-tubulin antibody (AB) and anti-FMRFa AB. The use of two neuronal markers allows visualization of a much larger spectrum of regulatory elements in the colony, enables separation of distinct neural subpopulations, and facilitates identification of novel conductive pathways for future analysis of the cellular basis of behavioral integration in these unique superorganisms.

**Figure 1.**
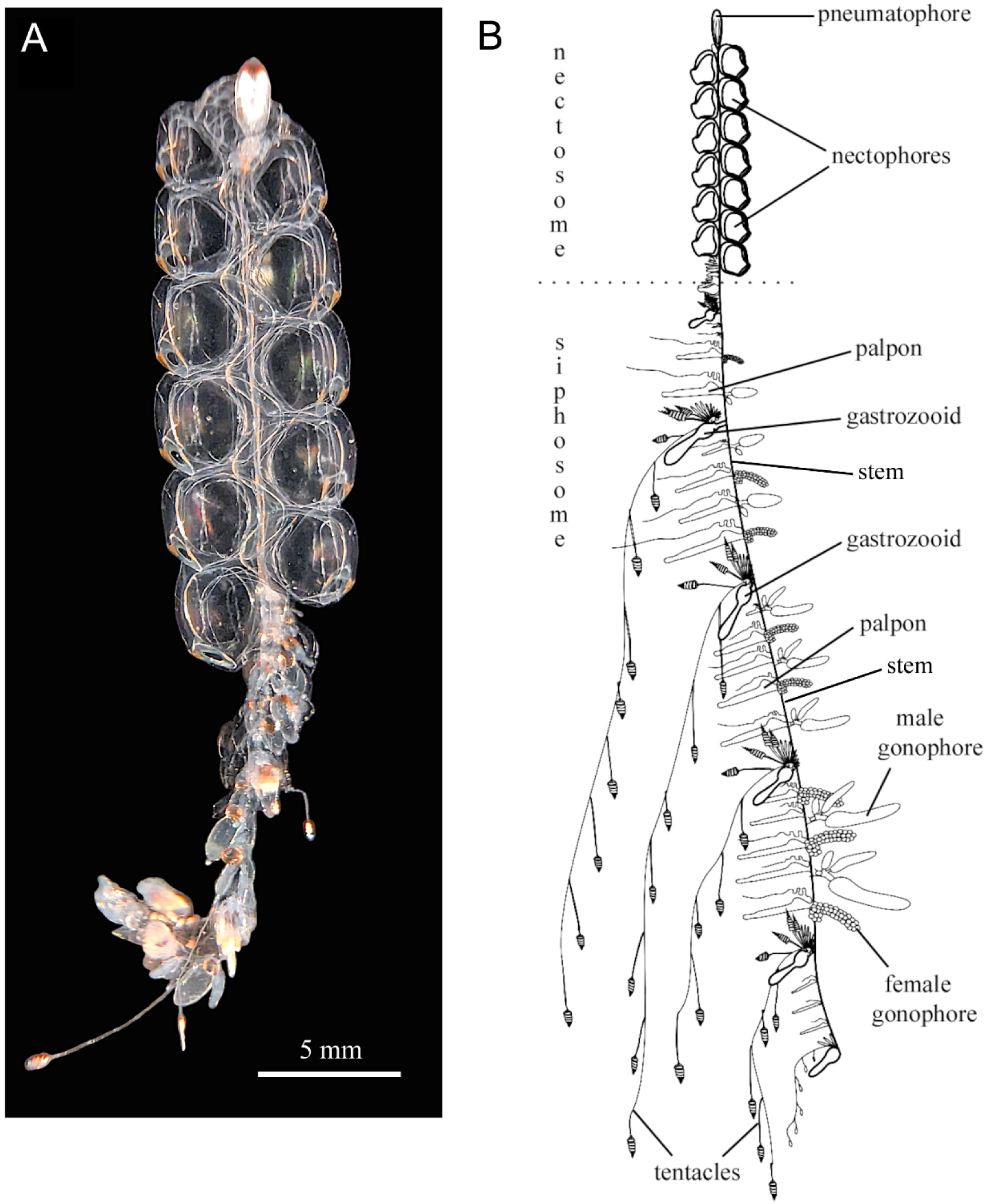
General view of *Nanomia septata* (formerly identified as *Nanomia bijuga*). (A) - Photo of live colony. (B) - Schematic drawing of the *Nanomia septata* with most zooids represented except for defensive bracts (for clarity reason). Modified from F. Goetz (https://commons.wikimedia.org/wiki/File:Nanomia_bijuga_whole_animal_and_growth_ zones.svg).

## MATERIALS AND METHODS

### Animals

Adult specimens of *Nanomia septata* (formerly identified as *Nanomia bijuga*, but later updated by Hosia et al., 2024) were collected from the dock at Friday Harbor Laboratories, University of Washington, USA, in the spring-summer and winter seasons of 2020-2025 and held in 1-gallon glass jars in large tanks with constantly circulating seawater at 10° C. Animals were fixed immediately after collection or survived in good condition for several days before being fixed.

### Immunocytochemistry

To keep the tissue relaxed in its natural form during fixation we incubated live animals in high Mg^2+^ solution (1 part 0.3M MgCl_2_ and 2 parts of filtered seawater) for 20 minutes. Immediately following that incubation, the animals were placed in the fixative - 4% paraformaldehyde in 0.1 M phosphate-buffered saline (PBS), overnight at +4-5° C. The fixed animals were then washed for several hours in PBS and dissected into several manageable pieces, usually separating the nectosome region and cutting the siphosome region into several sections, sometimes isolating individual gastrozooids.

To visualize the nervous system, we used two well-known markers. The first was anti-tubulin antibody (rat monoclonal antibody; AbD Serotec, Bio-Rad, Cat# MCA77G, RRID: AB_325003; recognizes the alpha subunit of α-tubulin, specifically binding tyrosylated α-tubulin (Wehland and Willingham, 1983, Wehland et al., 1983). We have successfully used this anti-tubulin antibody to label the neural systems in *Nanomia* nectophores and stem (Norekian and Meech, 2020, Norekian and Meech, 2026), in the hydrozoan *Aglantha digitale* (Norekian and Moroz, 2020a) and several ctenophore species (Norekian and Moroz, 2016, Norekian and Moroz, 2019a, Norekian and Moroz, 2019b, Norekian and Moroz, 2020b, Norekian and Moroz, 2021). The second marker was the anti-FMRFamide antibody (rabbit polyclonal; Millipore, Sigma, Cat# AB15348, RRID: AB_805291). The anti-FMRFamide antibodies have been shown to label the entire family of RFamide neuropeptides, which are all known to be present in a diversity of neural systems across Metazoa (Greenberg, 1988, Grimmelikhuijzen and Spencer, 1984, Mackie et al., 1985, Satterlie, 2008, Satterlie, 2014). Thus, anti-FMRFamide antibodies were used not to locate specifically FMRFamide itself, but as an additional marker of the neural (peptidergic) elements in *Nanomia*. We have successfully used this anti-FMRFamide antibody and anti-tubulin antibody combination before as two complementary neuronal markers in *Nanomia* nectosomal stem (Norekian and Meech, 2026) and in the hydrozoan *Aglantha digitale* (Norekian and Moroz, 2020a).

The dissected tissues were pre-incubated overnight in a blocking solution of 6% goat serum in PBS with 0.2% Triton-X and then incubated for 48 hours at +4-5° C in the primary antibodies. Anti-tubulin antibodies were diluted in 6% of the goat serum at 1:100, while anti-FMRFamide antibodies were diluted in 6% goat serum at 1:200. Following the primary antibody incubation and several PBS rinses over 12 hours, the tissues were placed for 24 hours in the secondary antibodies - goat anti-rabbit IgG antibodies, Alexa Fluor 568 (Thermo-Fisher, Cat# A-11011, RRID: AB_143157), at a final dilution 1:80 for FMRFa antibodies. For anti-tubulin antibodies we used goat anti-rat IgG antibodies, Alexa Fluor 488 conjugated (Molecular Probes, Invitrogen, Cat# A11006, RRID: AB_141373), at a final dilution 1:80 in the same solution. After 12 hours of washing, the tissue was mounted on glass microscope slides.

To label the muscle fibers, we used well-known marker phalloidin (Alexa Fluor 568 phalloidin from Molecular Probes), which binds to F-actin (Wulf et al., 1979). The fixed tissues were incubated in phalloidin solution in PBS for 8 hours at a final dilution 1:100 and then washed in several PBS rinses for 12 hours. All tissues were mounted on glass microscope slides in Vectashield mounting medium containing DAPI for nuclear labeling. The preparations were viewed using a Nikon Eclipse E800 microscope with epi-fluorescence using standard TRITC and FITC filters, and a Nikon C1 Laser Scanning confocal microscope. Over the course of 6 years of this investigation, we processed about 150 *Nanomia* with tubulin IR and phalloidin labeling, making more than 500 confocal scans, and about 100 *Nanomia* double-labeled with tubulin IR and FMRFa IR, taking about 500 confocal images (total 250 animals and more than a thousand confocal images). We studied mature animals/zooids (no developmental stages), which were fixed within 5 days of captivity.

## RESULTS

*Nanomia*’s extended stem runs through the entire length of the colony and serves as an anchor for numerous specialized zooids (Fig. 1). At the very anterior end, the stem connects to the float (pneumatophore) with a gas chamber, which is filled mostly with carbon monoxide (Pickwell et al., 1964). Attached to the stem, just under the float, are the medusa-like swimming zooids or nectophores, that propel the colony. The anterior part of the siphonophore colony, which includes two rows of nectophores, is called the nectosome (Fig. 1). New nectophores are constantly added from the growth zone, a region at the boundary between the pneumatophore and nectophores (Siebert et al., 2015). Below the nectosome, there is the siphosome region containing the colony’s other specialized zooids (Fig. 1). It includes gastrozooids, specialized for prey capture via outspread tentacles and initial digestion; palpons specialized for digestion only; bracts that serve a protective function; and reproductive gonads (male and female gonophores). New zooids are added from the second growth zone, at the anterior part of the siphosome (Siebert et al., 2015).

### Structure of the nervous system in Nectophores

Figure 2 shows the pneumatophore and nectophores in live *Nanomia*. Each nectophore’s main nerve complex consists of a ring nerve around the velum, laterally symmetrical column-shaped matrices, and two nerves (Fig. 3A, B; see more details about the nectophore organization, including electron microscopy and electrophysiology in Norekian and Meech, 2020). The nectophore ring nerve structure and function appear similar to those of other hydrozoan medusae (Satterlie, 2008). The lateral column-shaped matrices are unusual and may indicate additional information processing (Fig. 3B, C). The extensively branching upper nerve consists of many neural processes and appears to have a sensory role, collecting information from the large upper areas and directing it to the ring nerve (Fig. 3A, D). The lower nerve provides motor input/output, connecting the nectophore with the stem and the rest of the colony (Fig. 3A, B, E). The thin, unbranching lower nerve can be traced to the point of contact between the nectophore and the stem, where it ends in the terminal ganglion (Fig. 3E), which consists of 40–50 tightly packed neural cells (Norekian and Meech, 2020). This terminal ganglion is likely the most important element responsible for communication between the nectophore and the stem, and, therefore, the rest of the colony (Norekian and Meech, 2026).

**Figure 2.**
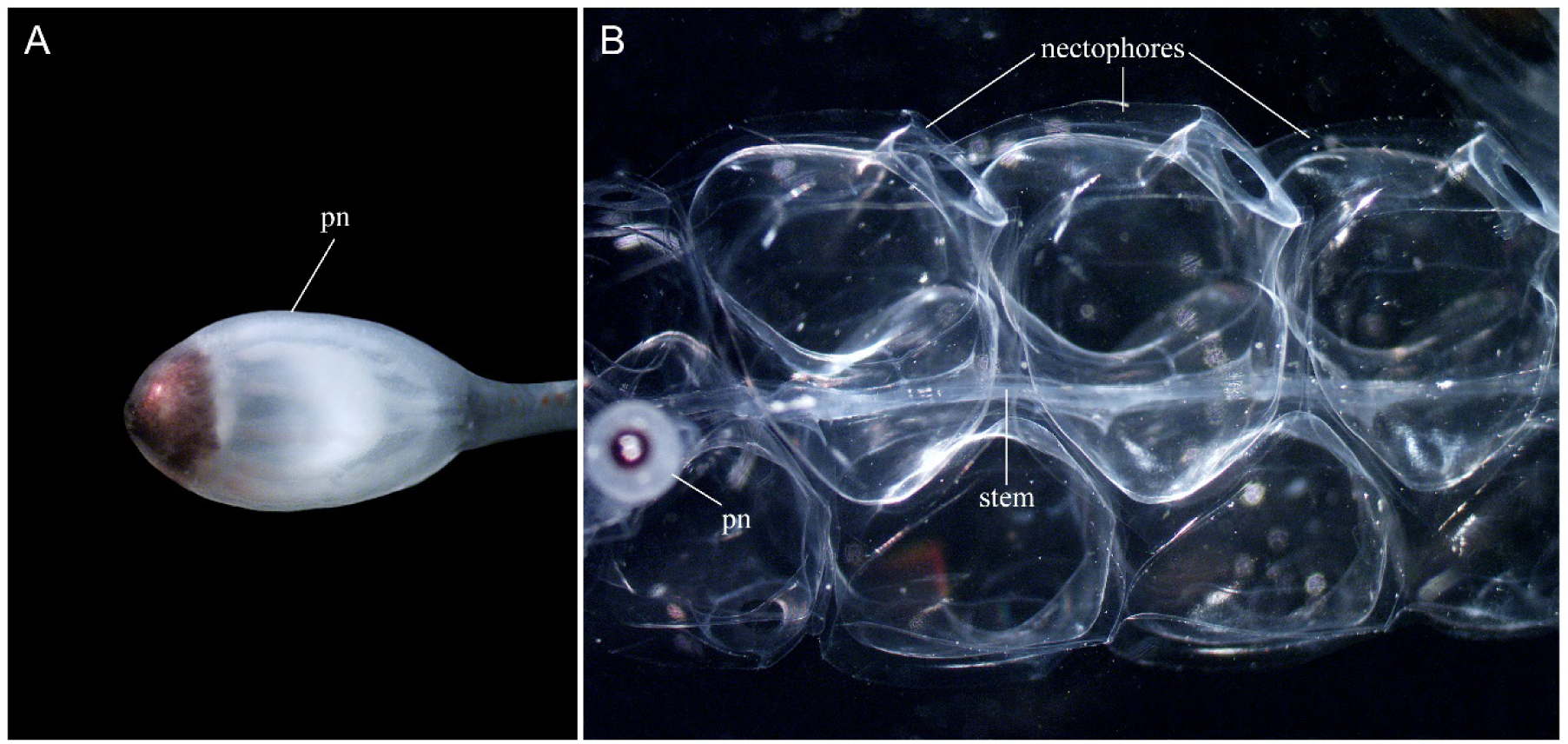
Light microscopy images of the anterior part of live *Nanomia*, including a pneumatophore (pn) - a gas-filled “float” (A) and the nectosome area with two rows of propulsive zooids - nectophores, which serve a locomotory function and resemble small medusa bells attached to the central stem (B).

**Figure 3.**
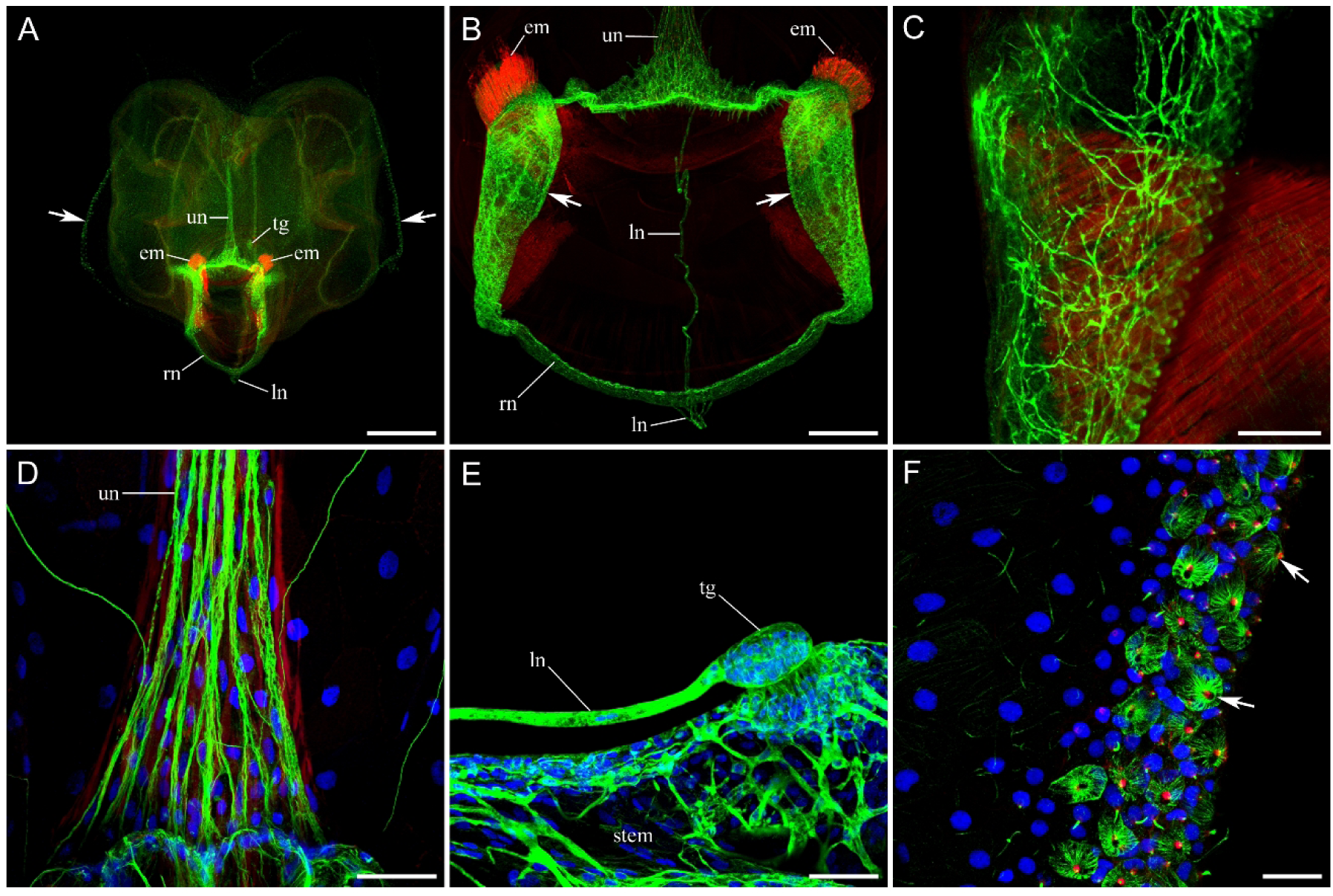
The neural system in nectophores is labeled with anti-tubulin AB (green). Muscles are stained with phalloidin (red), and cell nuclei are labeled with DAPI (blue). (A) - Image of an entire single nectophore. The nervous system contains the ring nerve (rn) around the velum and two nerves: the upper nerve (un) and the lower nerve (ln). The lower nerve (ln) ends in the terminal ganglion (tg) at the point of connection between the nectophore and the stem. Arrows point to the side ridges that contain numerous stinging cells - nematocytes. Two symmetrical bundles of endodermal muscles (em) are seen next to the velum. (B) - Higher magnification of the ring nerve area. A wide delta-shaped base connects the upper nerve (un) with the ring nerve (rn) at the top of the image. On the other side, a thin, non-branching lower nerve (ln) merges with the ring nerve (rn). Arrows point to the two laterally symmetrical, column-shaped matrices adjacent to the ring nerve. (C) - Higher magnification of the neural plexus in one of the columns. (D) - Higher magnification of the delta-shaped base of the upper nerve (un) next to the ring nerve. (E) - The lower nerve (ln) ends in the terminal ganglion (tg) at the point of connection between the nectophore and the stem. The terminal ganglion (tg) of each nectophore is separated from the stem surface by only a narrow cleft. On the other side of this cleft, in the stem, numerous tubulin-ir neural fibers also approach the surface. (F) - Numerous cone-shaped nematocytes (some are shown by arrows) at the lateral ridges of a nectophore, which are indicated in part (A). Scale bars: A - 500 µm; B - 200 µm; C - 20 µm; D - 40 µm; E - 50 µm; F - 20 µm.

On the stem side, there is a cone-shaped structure that serves as a nectophore docking station and is densely innervated by a tubulin-ir polygonal diffuse neural network (Fig. 3E; Norekian and Meech, 2026). The ectodermal epithelial layer that covers the nectophore’s outer surface is absent around the terminal ganglion, leaving only a narrow cleft between the ganglion and the cone surface (Fig. 3E; Norekian and Meech, 2026). Such morphological structure of the nectophore attachment area suggests the existence of a synaptic-like connection between the nectophore terminal ganglion and the diffuse neural network in the stem cone, including the possibility of volume transmission.

Although the upper nerve, the lower nerve, and the terminal ganglion in *Nanomia* nectophores do not show FMRFa immunoreactivity, the ring nerve and adjacent projections are rich in FMRFa-ir processes and neural cell bodies. Most fibers of the ring nerve are tubulin-ir only, yet several very thin FMRFa-ir processes contribute to the entirety of the ring nerve (Fig. 4A). Multiple FMRFa-ir neural cell bodies are also found adjacent to the ring nerve (white arrows in Fig. 4A; see also Grimmelikhuijzen et al., 1986). Many are located in the upper region of the velum on both sides of the upper nerve. The narrow lateral neural networks on the ring nerve and two neural projections extending from them (Norekian and Meech, 2020) also contain numerous thin FMRFa-ir processes (Fig. 4B; see also Grimmelikhuijzen et al., 1986). These include the shorter ‘z’ projection, directed outward toward the bell surface to innervate the epithelial ‘seitliche Zapfen’ area (Claus, 1878), and the second long cone-shaped projection extending toward the velum and Claus’ muscles. Some of these FMRFa-ir processes do not show double-labeling with tubulin IR, while others do.

**Figure 4.**
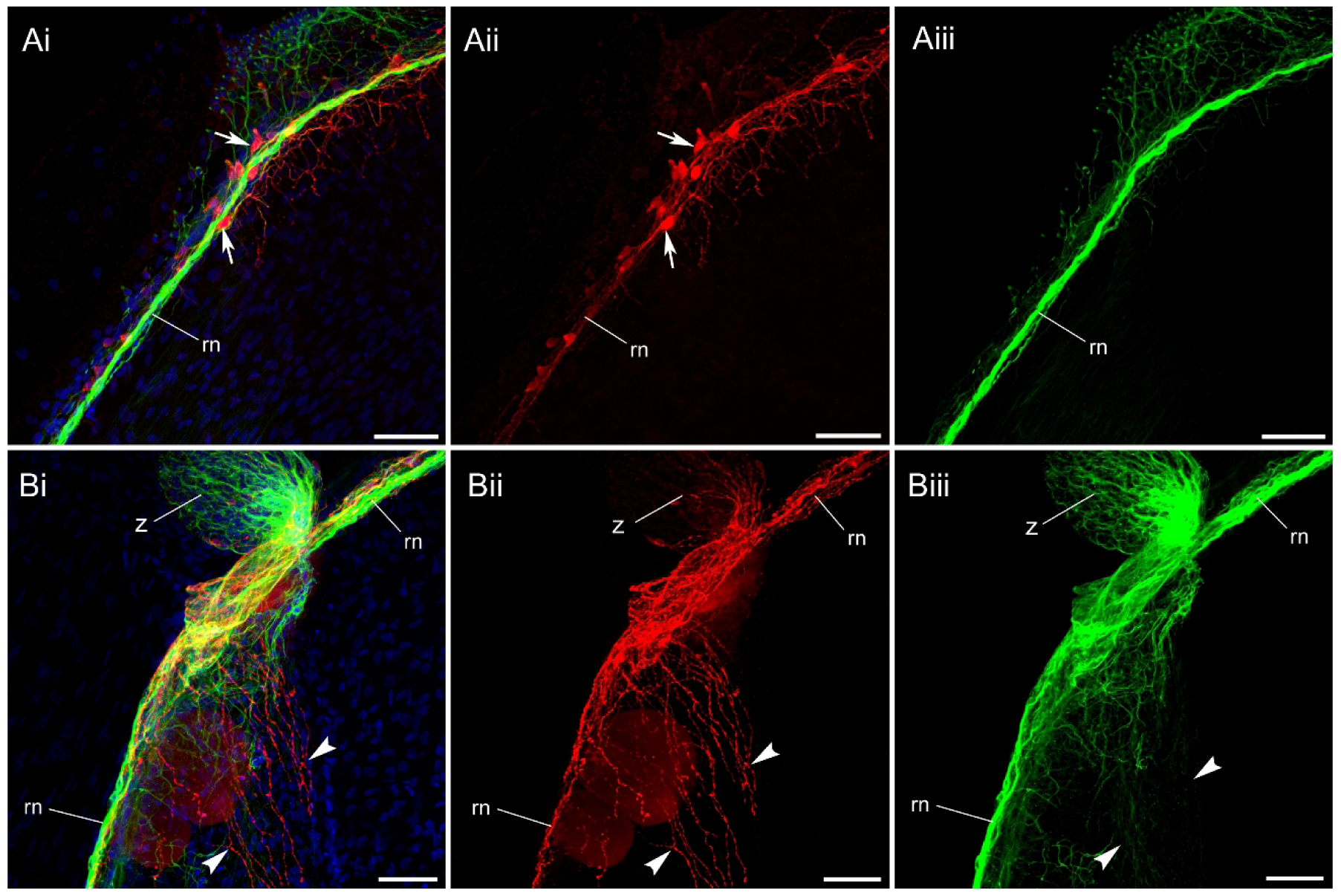
FMRFa-ir neural system in the nectophore ring nerve. Tubulin-ir labeling is green, FMRFa-ir labeling is red, and DAPI staining is blue. (Ai) - Double-labeling of the ring nerve (rn) near the upper nerve. Although the ring nerve is intensely labeled with anti-tubulin AB (green), it also contains neural cell bodies (arrows) labeled with anti-FMRF antibody (red). (Aii) - Red channel showing FMRFa-ir neurons and processes only. (Aiii) - Green channel with tubulin-ir only. (Bi) - Double-labeling preparation showing two projections connected to the ring nerve (rn): one projection (z) extends toward the ‘seitliche Zapfen’ area (Claus, 1878) in the epithelium, and the second (arrowheads) projects into the velum toward the Claus fibers. Both projections contain tubulin-ir and FMRFa-ir neural processes. (Bii) - Red channel with FMRFa-ir only. (Biii) - Green channel with tubulin-ir only. Scale bars: 50 µm.

### Structure of the nervous system in the Stem

Tubulin immunoreactivity revealed two main elements of the neural system in the stem in the nectosome area (Norekian and Meech, 2026). The first is a giant axon system, and the second is a polygonal subepithelial neural network. The giant axon runs from the anterior end of the stem at the base of the pneumatophore, through the entire length of the nectosome region, and then into the siphosome (Fig. 5A). In many preparations, we observed not one but two giant axons, located symmetrically on opposite sides of the stem, each about 40-50 μm in diameter (Fig. 6A; Norekian and Meech, 2026). The *Nanomia* nectosome area has two columns of nectophores, and presumably each axon in the stem is associated with one neighboring nectophore column. The stem also contains a diffuse polygonal neural network, located in the ectoderm and covering the entire stem surface at its entire length (Fig. 5B and 6A-C; Norekian and Meech, 2026). The individual units of the polygonal network vary in shape from triangles to octagons and in size from 10 to 100 μm (Fig. 6C). A notable feature of the stem in the nectosome area is the presence of multiple cone-shaped protrusions, which serve as docking sites for nectophores (Fig. 5B, C and 6A, B). Mackie described them as “muscular pedicle” (Mackie, 1964), while Grimmelikhuijzen et al. (1986) identified them as anchors or muscular lamellae. There are two columns of evenly spaced cones in the stem, corresponding to the two columns of nectophores. The stem polygonal neural network covers the entire cone protrusion from its base to the top surface where the nectophore is attached (Fig. 5B, C, and 6A, B; Norekian and Meech, 2026), thus providing the basis for neural connection between the nectophore and the central stem.

**Figure 5.**
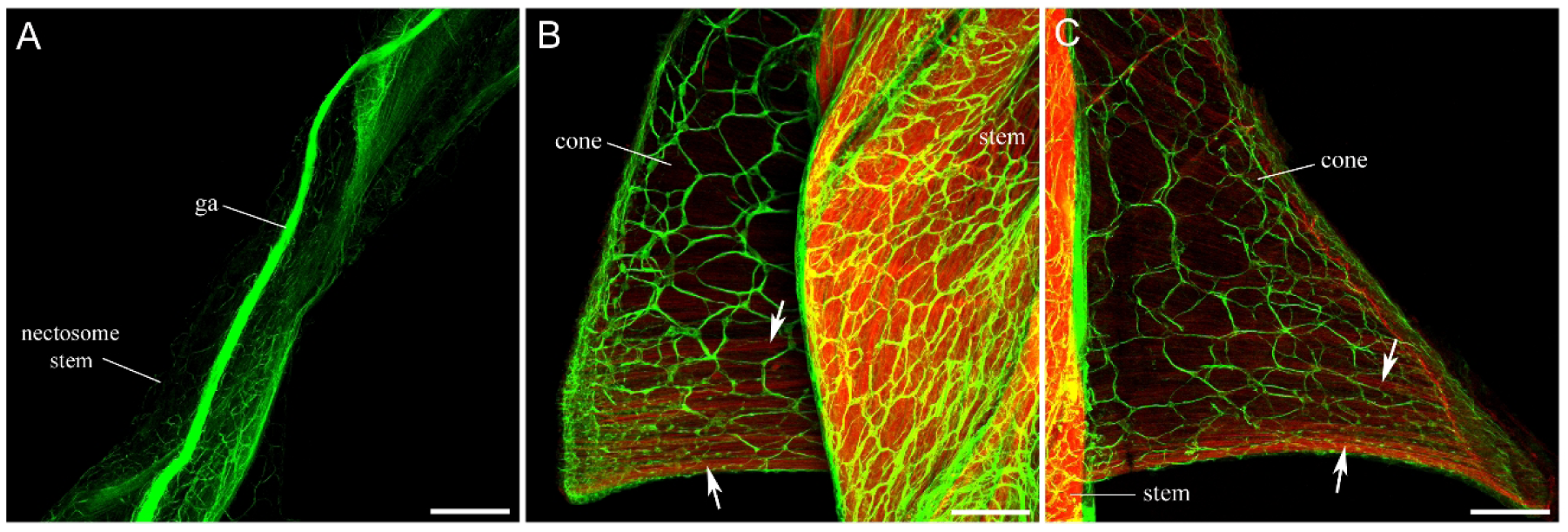
Neural system in the nectosome region of the stem, labeled with anti-tubulin antibody (green). Muscle system is labeled with phalloidin (red). (A) - Giant axon (ga) runs longitudinally through the entire length of the nectosomal stem. (B, C) - Tubulin-ir polygonal neural network covers the entire stem, including the cone-shaped protrusions that serve as docking sites for nectophores (cones). The stem itself is very muscular and brightly labeled with phalloidin. There are also muscle fibers (arrows) in the cones, suggesting their contractility. Scale bars: A - 200 µm; B, C - 100 µm.

**Figure 6.**
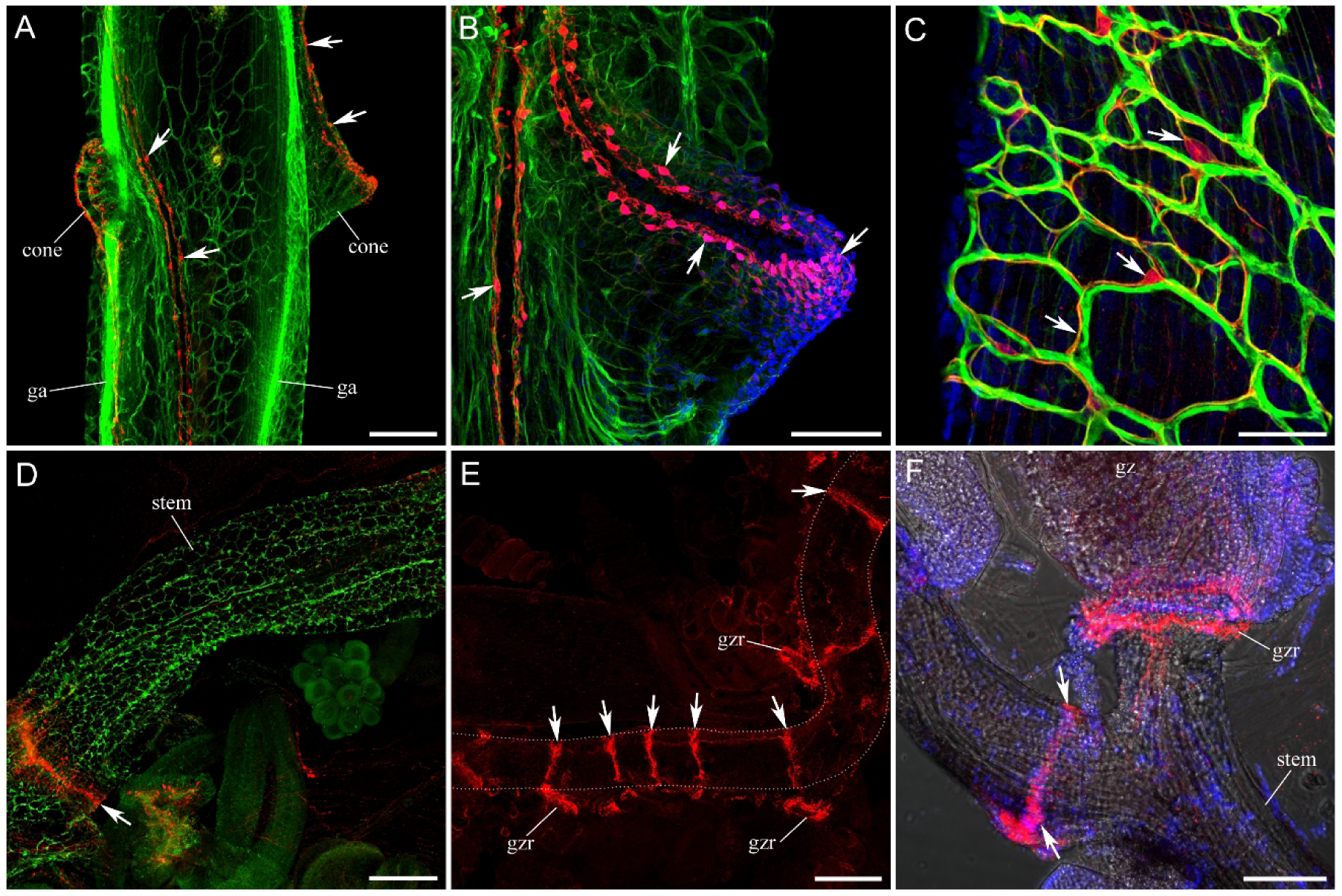
Structure of the nervous system in the stem, labeled with anti-tubulin antibody (green) and anti-FMRFa antibody (red). Nuclear DAPI staining is blue. (A) - Two giant axons (ga) run longitudinally through the stem in the nectosome region, one for each row of nectophores. Polygonal neural network covers the entire outer layer of the stem, including cones. There are also periodic FMRFa-ir neural double-stranded tracts (arrows) connecting the contralateral cones and forming a loop on their outer surface. (B) - High magnification of a cone showing a tubulin-ir polygonal network (green) and an FMRFa-ir neural loop (red) at the tip of the cone. Arrows point to some of the numerous FMRFa-ir cell bodies that form the tract. (C) - Double-labeling of the polygonal neural network in the nectosome stem revealed two structural subunits: one subunit has thicker threads and shows tubulin IR labeling only, while the other subunit (indicated by arrows) shows double-labeling with FMRFa IR. (D) - A diffuse polygonal neural network labeled with anti-tubulin AB (green) covers the stem in the siphosome region the same way as in the nectosome area. However, FMRFa-ir double-stranded tracts so typical for the nectosome region are absent. Instead, there are FMRFa-ir rings (arrow). (E) - These regularly located FMRFa-ir rings (arrows) divide the entire siphosome stem into sections. (F) - Higher magnification of the siphosome stem area with FMRFa IR (red) and the confocal transmitted light detector. FMRFa IR reveals the immunoreactive ring on the stem (arrows), as well as the ring at the gastrozooid base (gzr). Scale bars: A - 200 µm; B - 100 µm; C - 50 µm; D, F - 200 µm; E - 500 µm.

The stem of *Nanomia* is highly muscular and contractile (Mackie, 1973). Phalloidin labeling reveals that the stem is packed with thick muscle fibers (Fig. 5B and 7B, C, F). Of additional interest are the muscle fibers in the stem cones, oriented perpendicular to the stem from the cone base to its tip (Fig. 5B, C). Their contraction would shorten the cone and could be a mechanism for the precise, rapid separation of individual nectophores (autotomy), for example, during damaging impact on the colony.

FMRFa IR revealed one additional neural element in the nectosomal stem. Each cone, which serves as a nectophore docking site, is connected to another cone on the contralateral side (not its immediate neighbor but the next in the column) by a double-threaded FMRFa-ir pathway (Fig. 6A; Norekian and Meech, 2026). Each of these two threads of the FMRFa-ir pathway consists of multiple thin neural processes and regularly spaced neural cell bodies (Fig. 6A). The FMRFa-ir threads form a loop at the top of each cone on the side that connects to a nectophore. The loop at each cone contains numerous thin processes and multiple small immunoreactive neural cell bodies (Fig. 6B). This FMRFa-ir two-threaded pathway in the stem demonstrates a very strong immunoreactivity signal and shows no double-labeling with tubulin IR.

The FMRFa IR is also found in the polygonal neural network in the stem (Fig. 6C; Norekian and Meech, 2026). This FMRFa IR signal from the polygonal network is not as strong as that from the two-threaded pathway connecting contralateral cones, but it is highly consistent. A significant part of the tubulin-ir polygonal network, including its thickest threads, is tubulin-ir only and shows no double-labeling with FMRFa IR (Fig. 6C). However, the second part, consisting mostly of thinner processes, shows both tubulin IR and FMRFa IR (Fig. 6C). Thus, there are two subunits of the polygonal neural network in the stem. An identical situation is observed in the cones - one part of the polygonal neural network, which includes mostly thick threads, is tubulin-ir only, while the other part shows double-labeling with FMRFa IR. The RFamide-ir ectodermal neural network in the stem was also described earlier using different antibodies (Grimmelikhuijzen et al., 1986).

In the siphosome region, the structure of the neural system in the stem is slightly different. The diffuse tubulin-ir polygonal neural network covers the entire stem, as in the nectosome area (Fig. 6D and 7B). However, FMRFa-ir double-threaded pathways that connect the opposite cones of the stem in the nectosome region are absent. Instead, regularly spaced FMRFa-ir transverse bands (or collars) divide the entire siphosomal stem into sections or **cormidia** - repetitive clusters of various specialized zooids for tasks such as feeding (gastrozooids), reproduction (gonophores), and protection (bracts) that make up the siphosome region (Fig. 6D-F; see also Church et al., 2015, Grimmelikhuijzen et al., 1986). These transverse bands, or collars, are usually located posterior to each gastrozooid (Fig. 6F) and consist of many FMRFa-ir neurons and a network of their processes (see Fig. 14B). Labeling with phalloidin reveals that longitudinal muscle fibers are interrupted at the sites of the transverse bands (Fig. 15; similar to prior observations by Grimmelikhuijzen et al., 1986). In addition to the stem collars, FMRFa-ir labeling reveals numerous rings at the base of gastrozooids (Fig. 6E, F), which will be discussed in more detail below.

In the siphosome region, giant axons are located on the dorsal side of the stem, opposite to the zooid attachment side (Fig. 7A-C). They run longitudinally through most of the stem length, extending to the point where the stem begins branching at the very posterior end of the colony. In the region where the nectosomal stem transitions into the siphosomal stem, a giant axon crosses the stem from the ventral side in the nectosome region to the dorsal side in the siphosome region (Fig. 7D-F). Some images suggest that two axons cross the stem in that area, later merging into a single giant axon on the dorsal side of the siphosomal stem (Fig. 7E).

**Figure 7.**
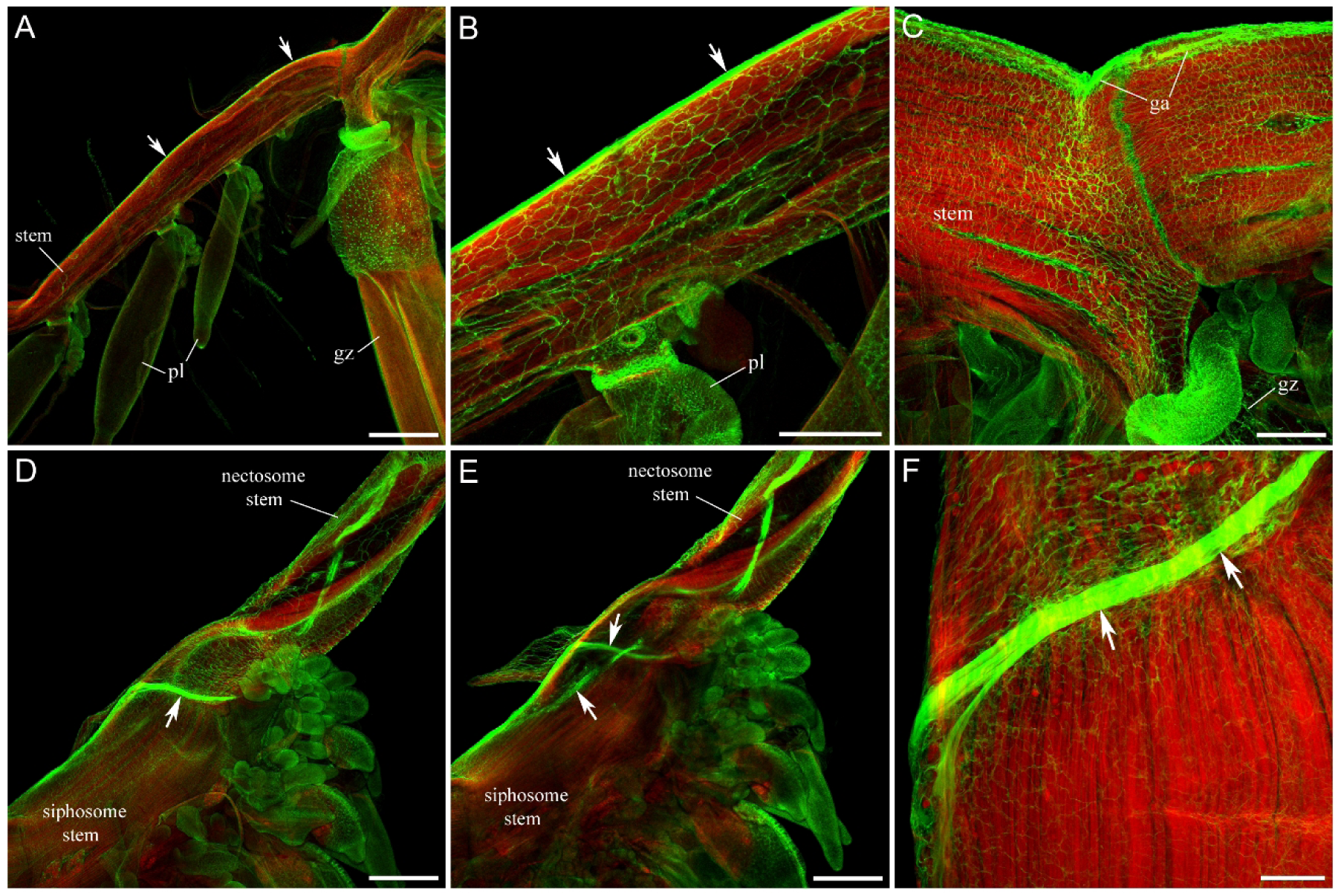
Giant axons in the siphosome region of the stem, labeled with anti-tubulin antibody (green). Phalloidin labeling of the muscles is red. (A) - Giant axon (arrows) runs longitudinally through the stem of the siphosome region on the dorsal side, opposite to the attachment side for other zooids. (B) - Higher magnification of the stem, with arrows pointing to the giant axon on the dorsal side, opposite to the palpon (pl) attachment. (C) - Giant axon (ga) runs along the dorsal side of the siphosomal stem, while a gastrozooid (gz) is attached to the ventral side of the stem via a short peduncle. Note the thick muscle fibers in the stem, labeled by phalloidin in red. (D) - This is the region where the nectosomal stem transitions into the siphosomal stem. Arrow points to the giant axon, which crosses the stem from the ventral side in the nectosome region to the dorsal side in the siphosome region. (E) - Optical section of the same preparation, deeper into the stem tissue, reveals two separate giant axons (arrows) that cross the stem and then merge into one in the siphosome region. (F) - High magnification of the giant axon (arrows) crossing the stem during the transition from nectosome to siphosome. Scale bars: A, D, E - 500 µm; B, C - 200 µm; F - 100 µm.

### Pneumatophore

The pneumatophore is a float with a large, gas-filled chamber at the anterior end of the *Nanomia* colony (Fig. 2A). The subepithelial polygonal neural network, labeled by tubulin IR, uniformly covers the entire pneumatophore surface (Fig. 8A, B). The network consists of polygonal units of various shapes and sizes and is very similar to the network described in the stem. At the point where the pneumatophore connects to the stem, there is a smooth, uninterrupted transition from the pneumatophore network to the stem network, suggesting full integration (Fig. 8A). The FMRFa-ir neural system is represented by a mash of numerous small cell bodies scattered throughout the entire pneumatophore body wall (Fig. 8C), with a higher concentration at the apex, around the apical pore (Fig. 8E; similar to those described earlier by Church et al., 2015, Grimmelikhuijzen et al., 1986). Only at the pneumatophore base does the FMRFa-ir neural system show a clear polygonal structure matching the tubulin-ir network (Fig. 8C, D). We think that this short FMRFa-ir polygonal neural network at the pneumatophore base is a short extension of the stem network and not a transverse collar, as was suggested by Grimmelikhuijzen et al. (1986). Double-labeling preparations show that the polygonal network at the base of the pneumatophore has two subunits (similar to the staining in the stem): one labeled with tubulin IR only (indicated by white arrows in Fig. 8D), which extends through the entire pneumatophore body, while the other subunit has FMRFa immunoreactivity, and is limited to the pneumatophore base. The pneumatophore body wall is muscular, with both longitudinal and circular fibers clearly labeled with phalloidin.

**Figure 8.**
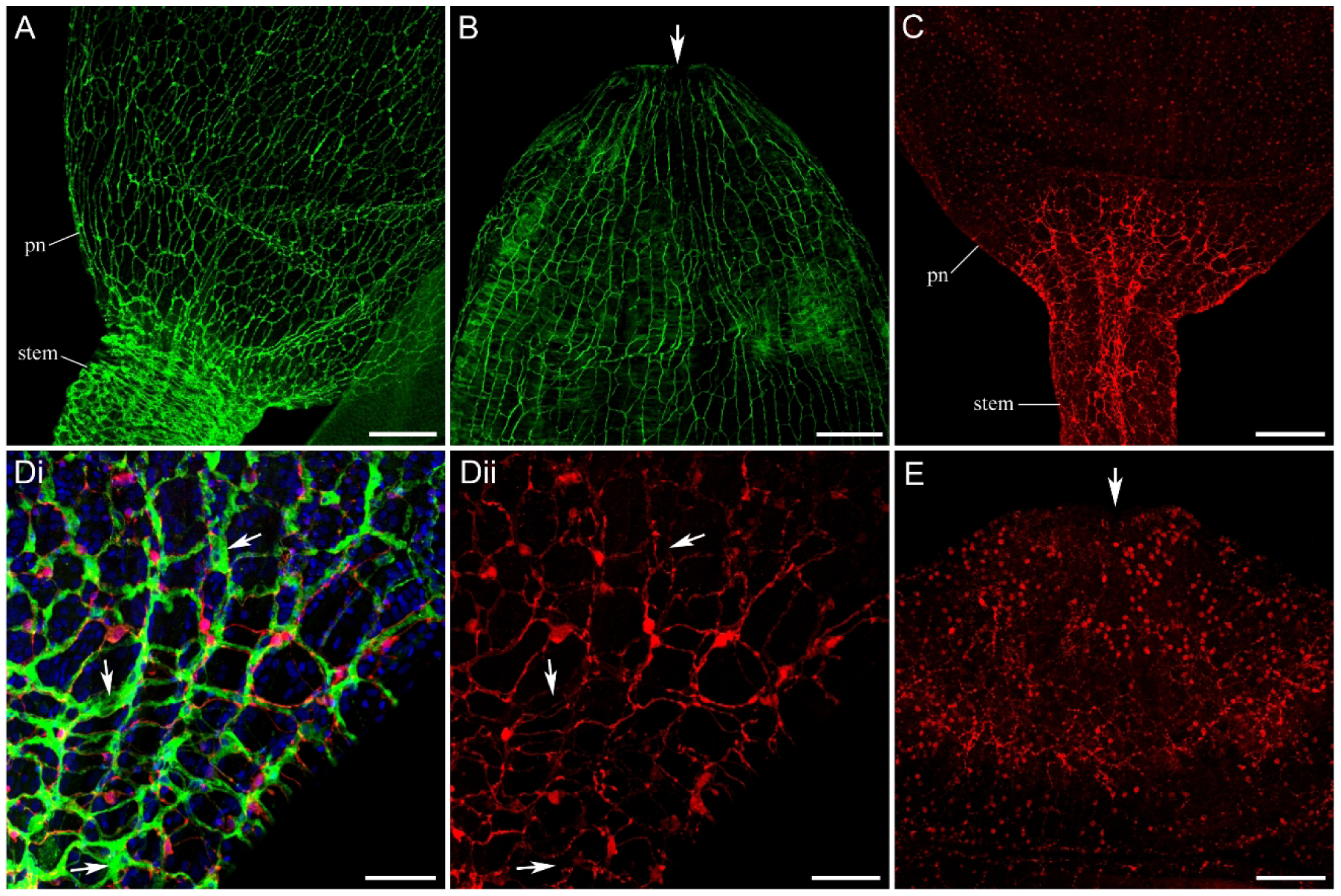
Neural network in the *Nanomia* pneumatophore (pn). (A) - Diffuse polygonal neural network labeled by tubulin IR (green) covers the entire outer layer of the pneumatophore and transitions without interruption to the stem. (B) - Tubulin-ir polygonal neural network at the apical side of the pneumatophore. Arrows point to the apical pore. (C) - FMRFa-ir neural system has a polygonal net structure only at the base of the pneumatophore, although numerous immunoreactive cell bodies are stained throughout the entire body wall. (Di) - Double-labeling at the base of the pneumatophore with tubulin IR (green) and FMRFa IR (red) shows that the polygonal neural network has two subunits: one labeled with tubulin IR only (indicated by white arrows), while the other subunit has FMRFa immunoreactivity - very similar to the staining in the stem. (Dii) - Red channel with FMRFa IR only. (E) - FMRFa-ir neurons in the apical area of the pneumatophore. The apical pore is shown by the arrow. Scale bars: A, B, C - 200 µm; D - 50 µm; E - 100 µm.

### Gastrozooids and palpons

Figure 9 shows the siphosome area in live *Nanomia* with the primary focus on gastrozooids and palpons. Each gastrozooid is a polyp specialized for feeding. It includes a short basigaster region, attached to the stem via a short peduncle, a much longer oral hypostome region, and a long tentacle with many branches called tentilla, which is attached to the base of a basigaster. There is also a morphologically distinct ring at the base of the basigaster (Fig. 9B-D). This ring is brightly stained with tubulin IR (Fig. 10A-C and 11A, B) and will be described below as a significant part of the nervous system in gastrozooids. The palpon is a polyp that is specialized for digestion but lacks the ability to capture prey and feed as gastrozooid. Mackie et al. (1988) described palpons as an accessory of the digestive system that can be seen inflating and deflating with gastric fluid. Palpons are noticeably smaller and narrower than gastrozooids. Phalloidin labeling of the muscle system also revealed that, while gastrozooids have a very muscular body wall with numerous longitudinal and circular muscle fibers, palpons do not have any (Fig. 10E, F).

**Figure 9.**
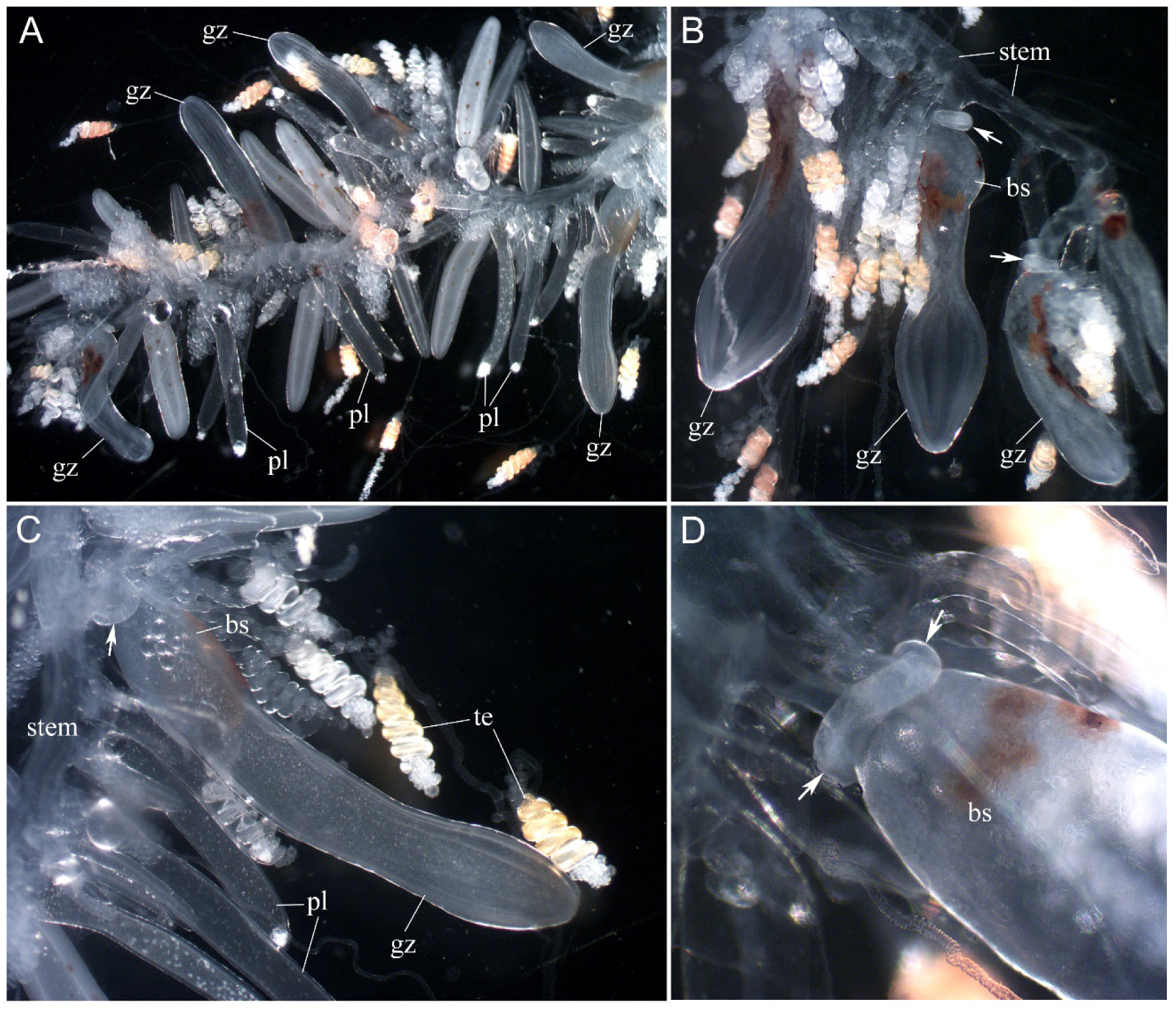
Light microscopy images of the siphosome area in live *Nanomia*, with a focus on gastrozooids (gz) and palpons (pl). (A) - Overview of the siphosome region at low magnification. (B) - Three gastrozooids (gz) attached to the stem at higher magnification. Note the ring at the base of each gastrozooid (arrows) attached to the basigaster (bs). (C) - Elongated single gastrozooid (gz) with tentilla (te) and several smaller palpons (pl). (D) - High magnification of the ring (arrows) at the base of a basigaster (bs).

**Figure 10.**
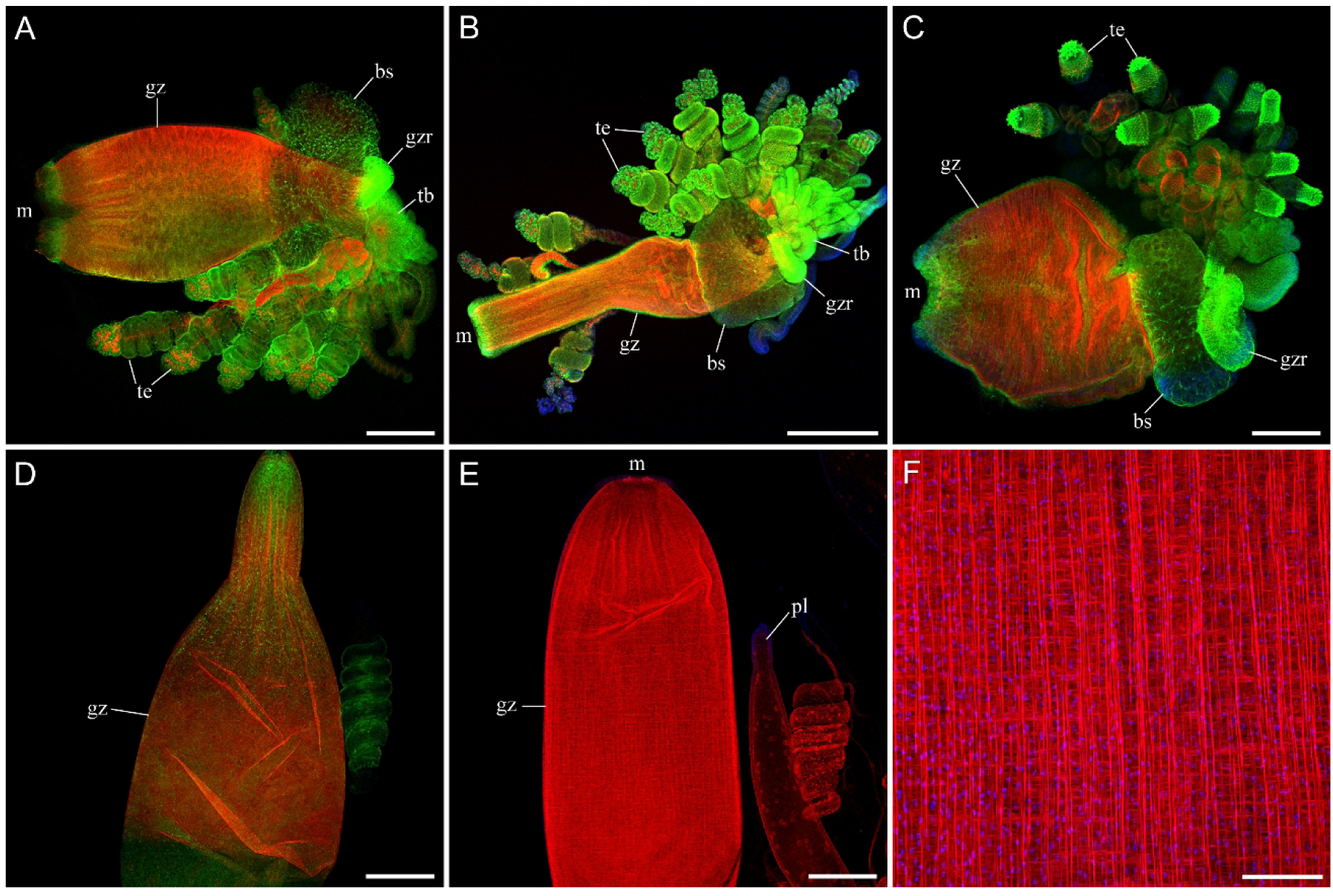
Overview of *Nanomia* gastrozooids of variable shapes and stages, outlined by phalloidin labeling in red and anti-tubulin AB in green (A-D). Note the basigaster (bs), or the base region; the hypostome region of gastrozooids (gz); mouth (m); tentacle base (tb); tentilla (te); and the gastrozooid ring (gzr) at the base, where it connects to the stem by a short branch. (E) - Phalloidin labeling (red), which visualizes the muscle fibers, shows that gastrozooid body wall is very muscular, in contrast to the palpon (pl) body wall. (F) - There are two types of muscle fibers in the gastrozooid wall: circular and longitudinal, which run perpendicular to each other. Scale bars: A-E - 500 µm; F - 100 µm.

In the basigaster region, tubulin IR reveals multiple branching neuronal-like processes (Fig. 11A-C). The entire elongated hypostome region is covered by a diffuse polygonal neural network (Fig. 11D, E). Some individual neurons with branching processes from that network are embedded within the muscular body wall of the hypostome among circular and longitudinal muscle fibers (Fig. 11F).

**Figure 11.**
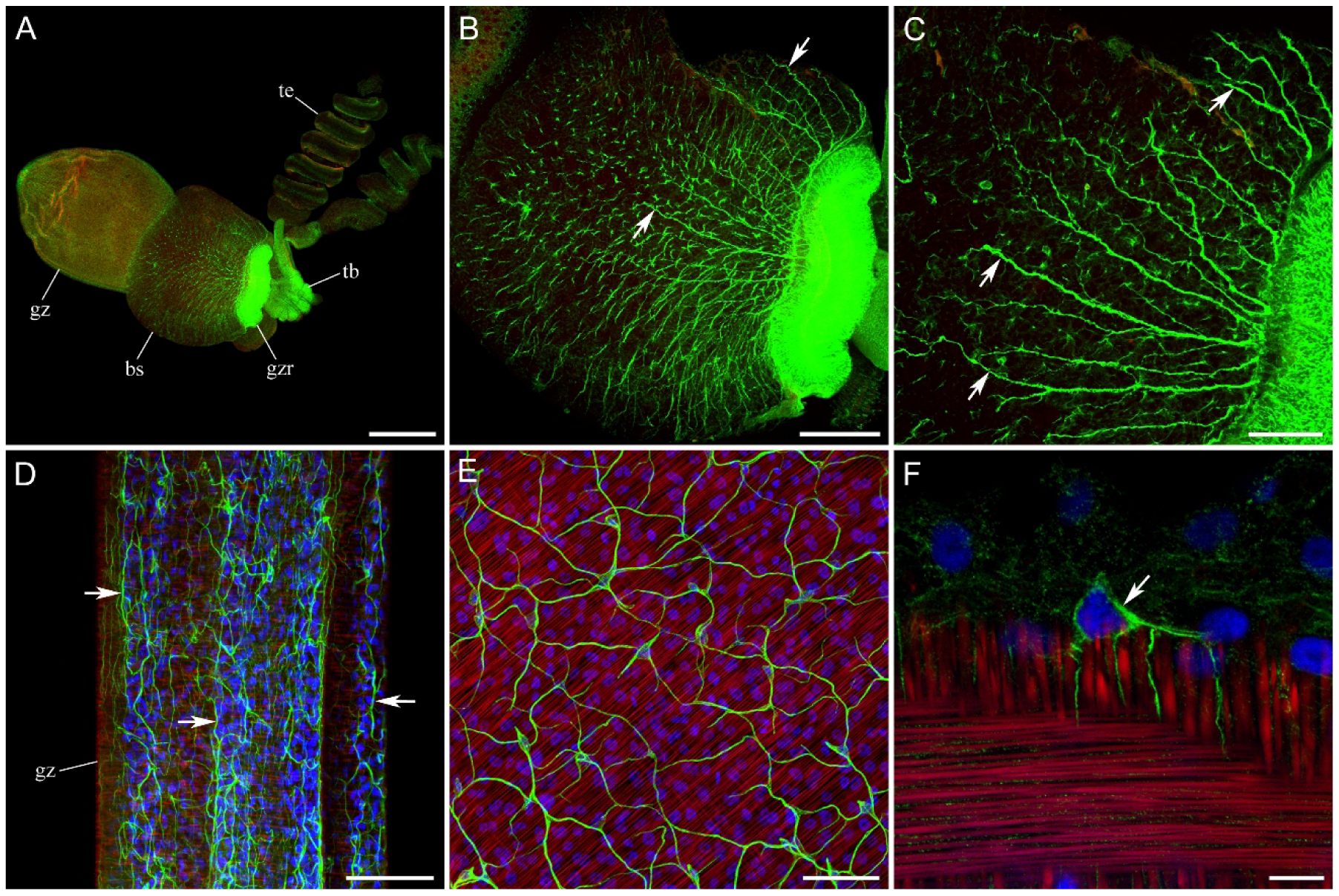
Neural network in gastrozooids labeled with anti-tubulin AB (green). Muscles are labeled with phalloidin in red, and nuclear DAPI is blue. (A-C) - In the basigaster (bs) region of gastrozooids, tubulin IR reveals multiple branching neuronal-like processes (some indicated by arrows). (D-E) - The entire hypostome region of each gastrozooid (gz) contains a diffuse polygonal neural network (arrows). (F) - An individual neuron (arrow) with several processes is embedded in the gastrozooid body wall among circular and longitudinal muscle fibers. The optical section is 10-15 µm thick. Other abbreviations: gzr - gastrozooid ring, tb - tentacle base, te - tentilla. Scale bars: A - 500 µm; B - 200 µm; C - 50 µm; D - 200 µm; E - 50 µm; F - 10 µm.

FMRFa IR revealed a very different population of neural elements (Fig. 12; comparable to prior observations (Church et al., 2015, Grimmelikhuijzen et al., 1986). The entire hypostome region contains many scattered FMRFa-ir neurons, associated with numerous unbranching, longitudinally running immunoreactive processes that extend from the basigaster region to the mouth (Fig. 12A-D). At the oral end of the hypostome, in the lip area around the mouth, there is a high density of presumably sensory FMRFa-ir neurons (Fig. 12D-F). These “sensory” cells are bipolar, with one long process running inward and a second short process or cilium projecting to the surface (Fig. 12F).

**Figure 12.**
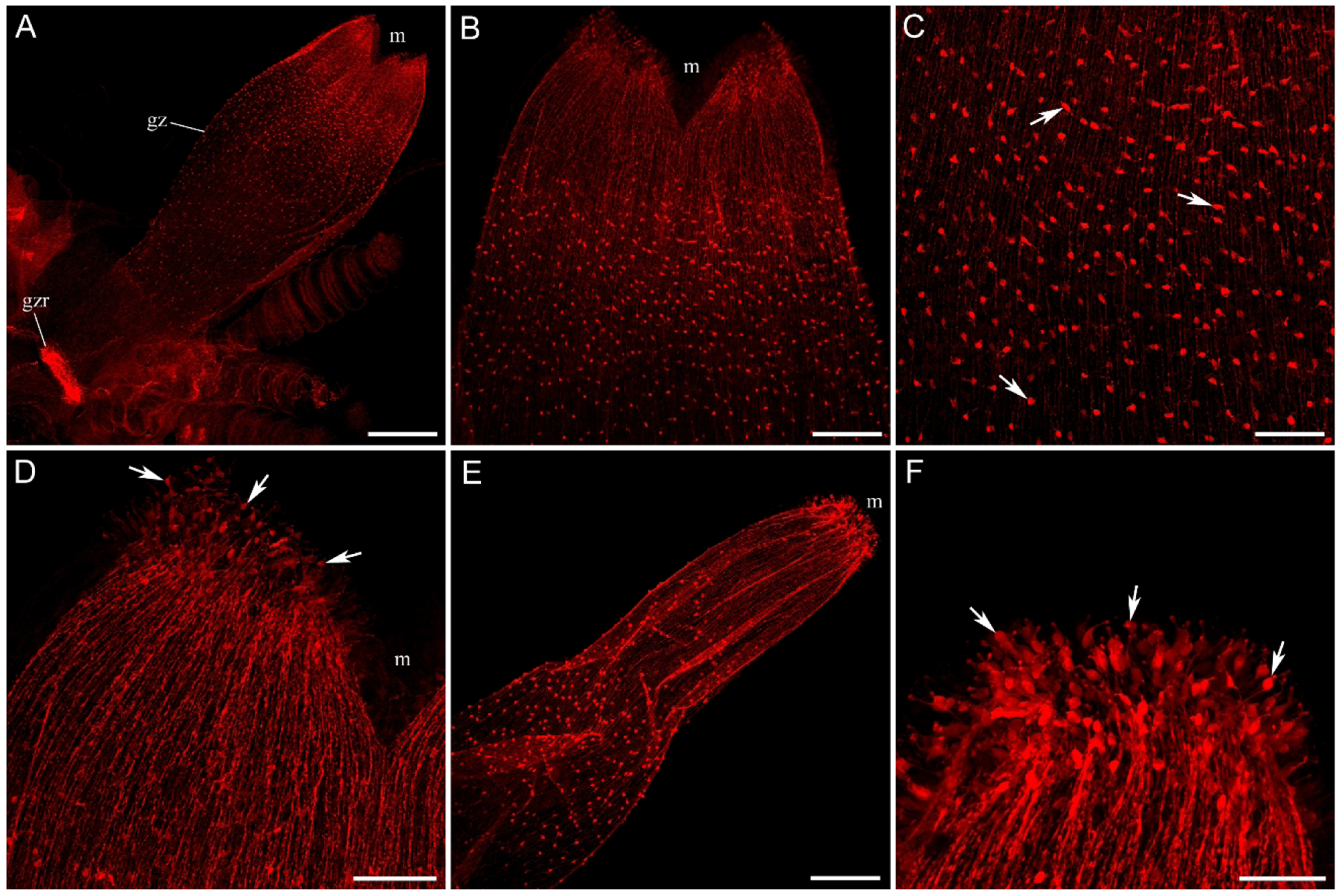
FMRFa-ir neural network in *Nanomia* gastrozooids. (A) - FMRFa-ir neural elements are found throughout the entire gastrozooid body, from the gastrozooid ring (gzr) at the base through the hypostome region to the buccal end (m - mouth). (B, C) - The entire hypostome region contains numerous small FMRFa-ir neuronal cell bodies (arrows), which produce longitudinally oriented parallel processes. (D) - At the buccal end, in the lip area, many presumably sensory neurons are labeled with FMRFa AB (arrows). (E) - Gastrozooids of all sizes and shapes have the same structure of the FMRFa-ir neural system. (F) - High magnification of the mouth area, with arrows pointing to some of the FMRFa-ir sensory neurons. Note that these cells are bipolar, with one long process running inward and a second short process projecting toward the outside surface. Scale bars: A - 500 µm; B - 200 µm; C-E - 100 µm; F - 50 µm.

Double-labeling of *Nanomia* gastrozooids with tubulin IR and FMRFa IR shows that there is no co-localization and that these two markers label completely separate subunits of the neural system in gastrozooids (Fig. 13). One is a polygonal neural network labeled by tubulin IR only. The other consists of small, scattered neurons with numerous parallel longitudinal processes and sensory neurons in the lip area, all labeled by FMRFa IR only. This is a clear example of how tubulin IR and FMRFa IR complement each other, revealing different populations of neural elements.

**Figure 13.**
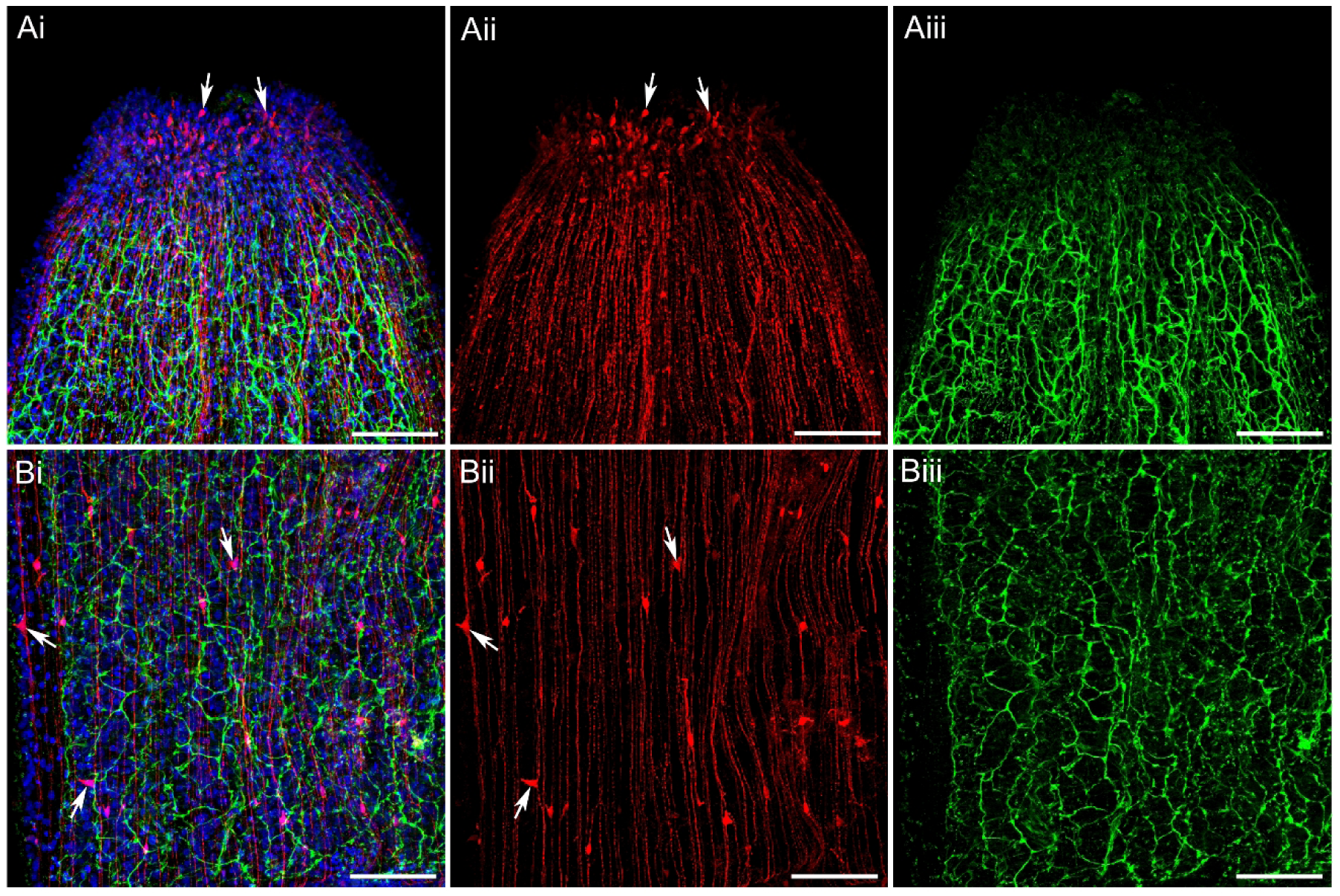
Double-labeling of *Nanomia* gastrozooids with tubulin IR (green) and FMRFa IR (red) reveals two separate neural subsystems. One is a polygonal neural network labeled only with tubulin IR (green). The other consists of numerous longitudinal processes running parallel to each other along the entire hypostome region and labeled only with FMRFa IR (red). They do not show any co-localization. (A) - Buccal hypostome region. Arrows point to numerous FMRFa-ir, presumably sensory neurons in the lip area around the mouth. (Ai) - Double-labeling, (Aii) - red channel showing FMRFa IR only, (Aiii) - green channel showing tubulin IR only. (B) - Mid-region hypostome area. Arrows point to some FMRFa-ir cell bodies that form parallel longitudinal processes. (Bi) - Double-labeling, (Bii) - red channel showing FMRFa IR only, (Biii) - green channel showing tubulin IR only. Scale bars: 100 µm.

One interesting morphological structure in gastrozooids is a ring at their base, which is visible under regular light microscopy (Fig. 9B-D) and also brightly stained with tubulin IR (Fig. 10A-C and 11A, B). This is also the location of densely packed FMRFa-ir neurons and processes that form a neural ring at the gastrozooid base (Fig. 12A and 14; see also Church et al., 2015, Grimmelikhuijzen et al., 1986). A bright tubulin-ir staining is probably not related to neural elements, but rather to the presence of densely packed, radially oriented tubulin filaments of non-neural origin (Fig. 14B, D). However, FMRFa IR clearly reveals numerous neural cell bodies and densely packed neural processes in the FMRFa-ir ring (Fig. 14B, D). Many of those neurons are bipolar and have a single short projection to the surface, suggesting they could be sensory cells (Fig. 14B, D). Longitudinally oriented FMRFa-ir neural processes run from the FMRFa-ir neural ring through the basigaster to the hypostome neural network (Fig. 14C). They also connect the FMRFa-ir ring with the stem neural network (Fig. 14B, C), thus integrating the gastrozooid and stem neural systems and coordinating the colony’s function.

**Figure 14.**
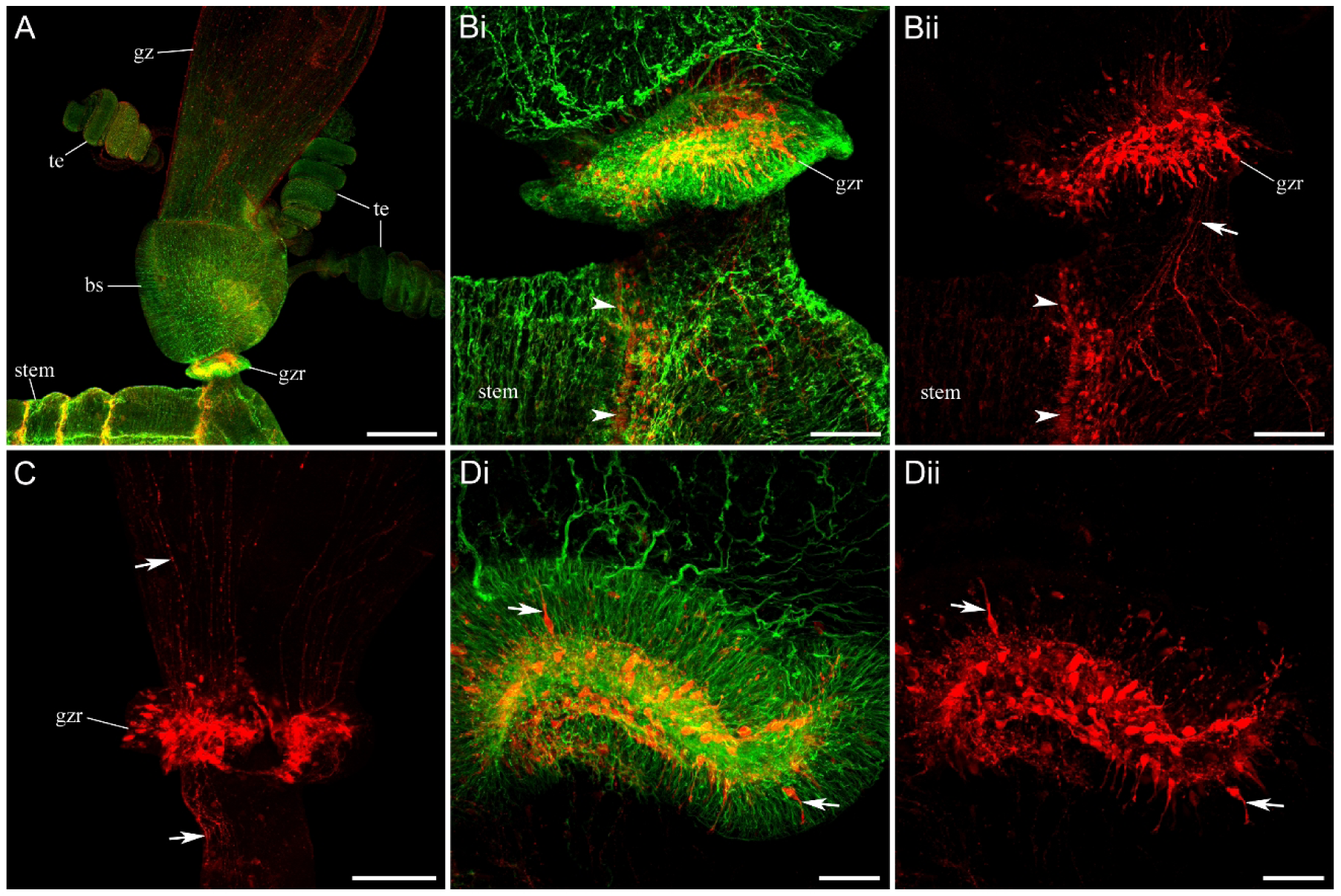
Nervous system in the gastrozooid ring, labeled with tubulin IR (green) and FMRFa IR (red). (A) - At the base of each gastrozooid (gz), below the basigaster (bs) region and above a short branch connecting gastrozooid to the stem, there is a gastrozooid ring (gzr) clearly visible morphologically (see also figure 7). (Bi) - This gastrozooid ring (gzr) is brightly labeled with tubulin AB (green) and contains numerous FMRFa-ir (red) neural cell bodies and their processes forming a ring around the central canal of the connecting branch. Arrowheads point to another FMRFa-ir ring in the stem. (Bii) - Red channel with FMRFa IR only. Arrow points to FMRFa-ir neural processes connecting the gastrozooid ring (gzr) with the stem neural network. (C) - Dense neural FMRFa-ir gastrozooid ring (gzr) integrates the FMRFa-ir neural systems of the gastrozooid hypostome and the stem: arrows point to the connecting processes. (D) - High-magnification of the gastrozooid ring, double-labeled with tubulin IR and FMRFa IR. Note that tubulin IR (green) labels numerous filaments in the ring, which might not have a neural origin. (Dii) - Red channel with FMRFa IR only. Arrows point to some of the FMRFa-ir neuronal cell bodies in the ring, with their specific morphology. Scale bars: A - 500 µm; B, C - 100 µm; D - 50 µm.

The gastric cavity of the stem and the gastric cavities of gastrozooids are connected and continuous. A hypothesis was proposed that the morphologically distinct ring at the gastrozooid base might function as a sphincter that contracts to isolate each gastrozooid from the stem. To function as a sphincter, it should have prominent circular muscles. However, phalloidin-labeling experiments demonstrated that gastrozooid rings contain no muscle fibers (Fig. 15), effectively ruling out their possible role as an isolating sphincter.

**Figure 15.**
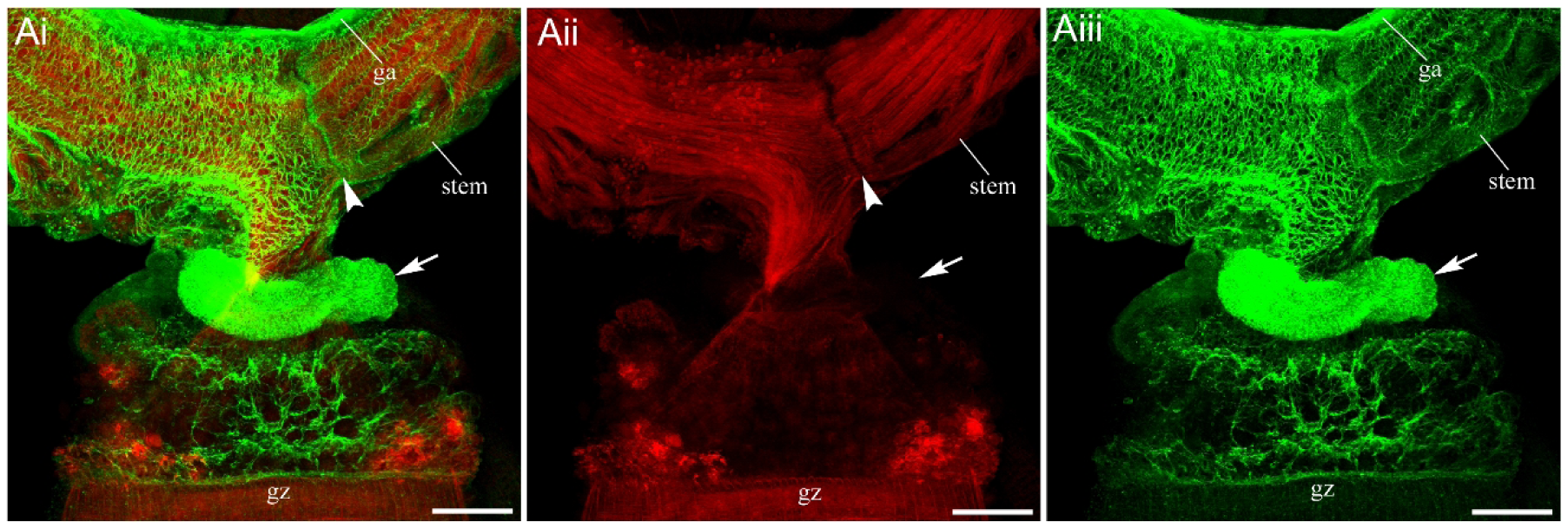
The gastrozooid ring lacks muscle fibers, as revealed by phalloidin (red) labeling. Tubulin IR is green. (Ai) - The gastrozooid ring (arrow) is clearly visible in tubulin IR (green) at the base of each gastrozooid (gz). (Aii) - Red channel with phalloidin labeling only. Note that while the stem contains many longitudinal muscle fibers throughout its entire thickness, the gastrozooid ring (arrow) lacks phalloidin labeling. Arrowhead points to the area where the longitudinal muscle fibers are interrupted and where the transverse FMRFa-ir band is located (not shown). (Aiii) - Green channel showing tubulin IR only. Abbreviations: ga - giant axon. Scale bars: A - 200 µm.

Each tentacle attached to the base of a gastrozooid has many side branches, called tentilla. Each tentillum is connected to the tentacle via a flexible, muscular pedicle (Figs. 16, and 17B). The spiraled cnidoband contains stinging cells, nematocysts that function as the primary prey-capture mechanism and are always brightly labeled by tubulin AB (Figs. 16B, and 17A,B). A terminal filament at the top sticks to a prey and then pulls the elastic strand, causing the cnidoband to fire nematocysts. The epithelial expansion called the involucrum extends from the pedicle and partially covers the cnidoband. FMRFa IR reveals a diffuse network of neurons and connecting processes on the surface of the cnidoband and in the terminal filament area (Fig. 18A, B). A much denser, well-developed neural network is observed in the pedicle and involucrum, as well as in the tentacle itself (Fig. 18C-E).

**Figure 16.**
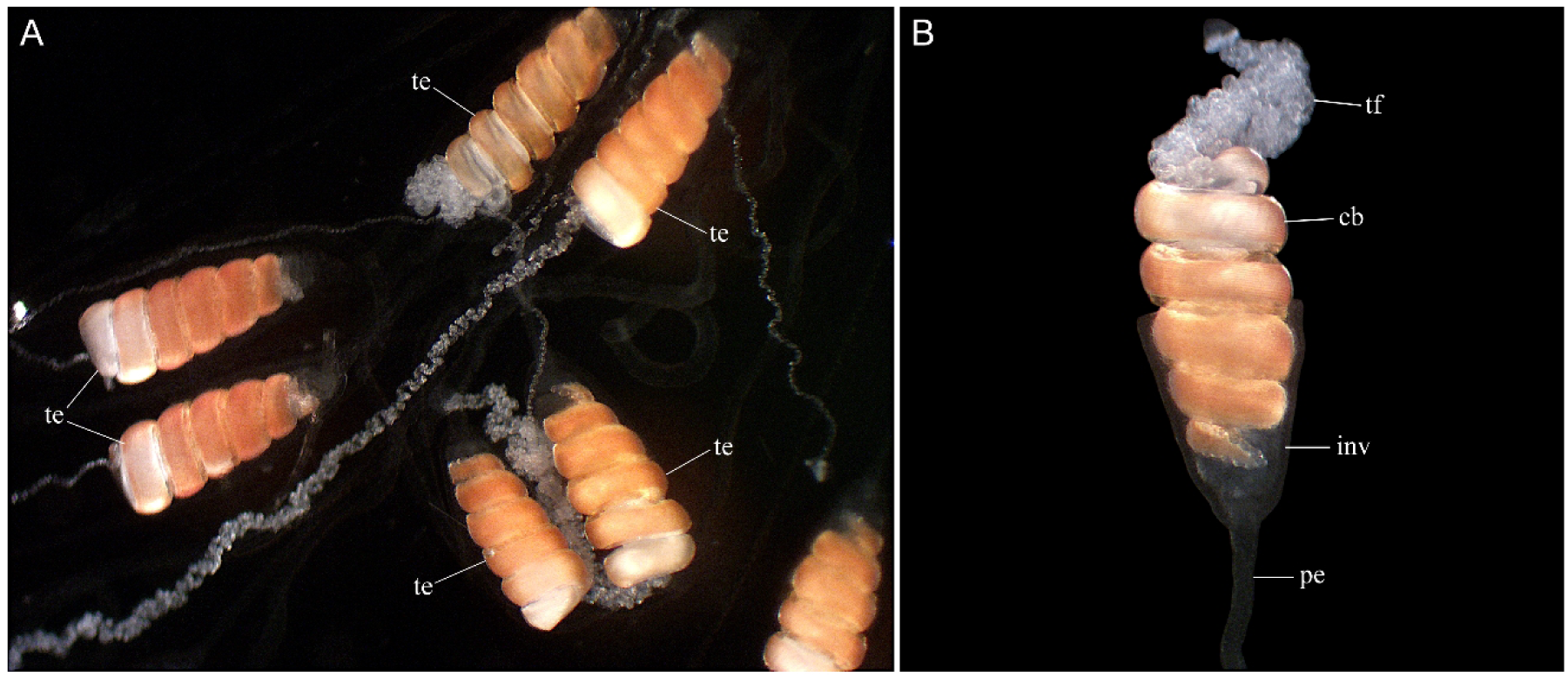
Light microscopy images of the tentilla. (A) Overview of the tentacle with several individual tentilla (te). (B) Morphological structure of a single tentillum. Each tentillum is connected to the tentacle via a flexible pedicle (pe). The spiraled cnidoband (cb) contains dense batteries of stinging cells (cnidocytes/nematocysts) that function as the primary prey-capture tool. A terminal filament (tf) at the top first sticks to a prey, pulling the elastic strand and causing the cnidoband to fire nematocysts. There is also an epithelial expansion called the involucrum (inv) that extends from the pedicle and partially covers the cnidoband.

**Figure 17.**
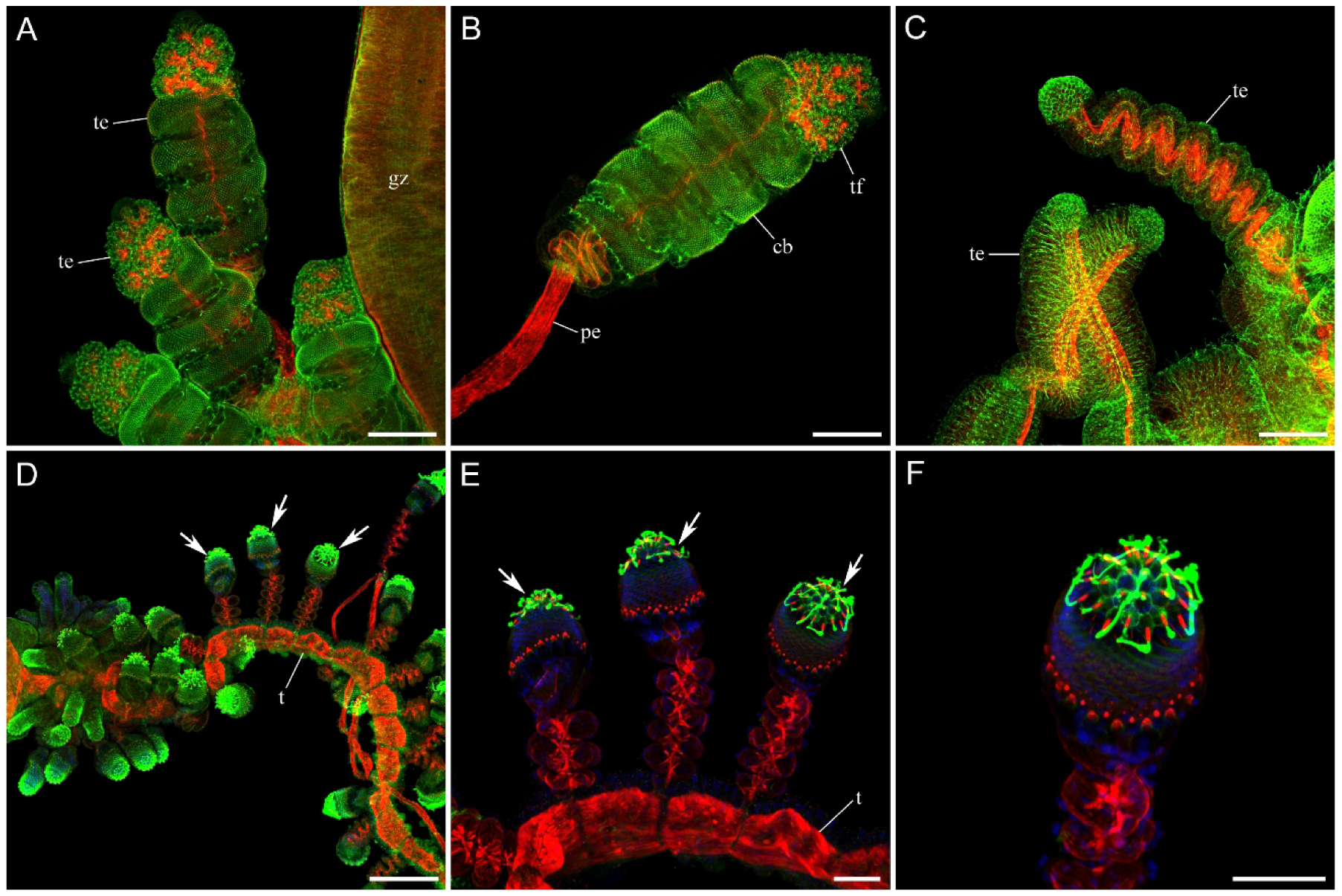
Overview of *Nanomia* tentacles at different stages, outlined by phalloidin labeling in red and anti-tubulin AB in green. (A) - Several mature tentilla (te) next to a gastrozooid (gz) body. (B) - A single mature tentillum with a muscular pedicle (pe) labeled by phalloidin, a spiraled cnidoband (cb) highlighted in green because tubulin AB target nematocysts, and the terminal filament (tf). (C) - A few tentilla (te) at their intermediate stage. Note the spiraled muscle fibers labeled with phalloidin. (D-F) - A young tentacle (t) with very small, undeveloped tentilla (arrows). Note that terminal filaments at this stage are represented by groups of individual long cilia, brightly labeled with tubulin IR (green). Scale bars: A, B - 200 µm; C - 50 µm; D - 200 µm; E, F - 50 µm.

**Figure 18.**
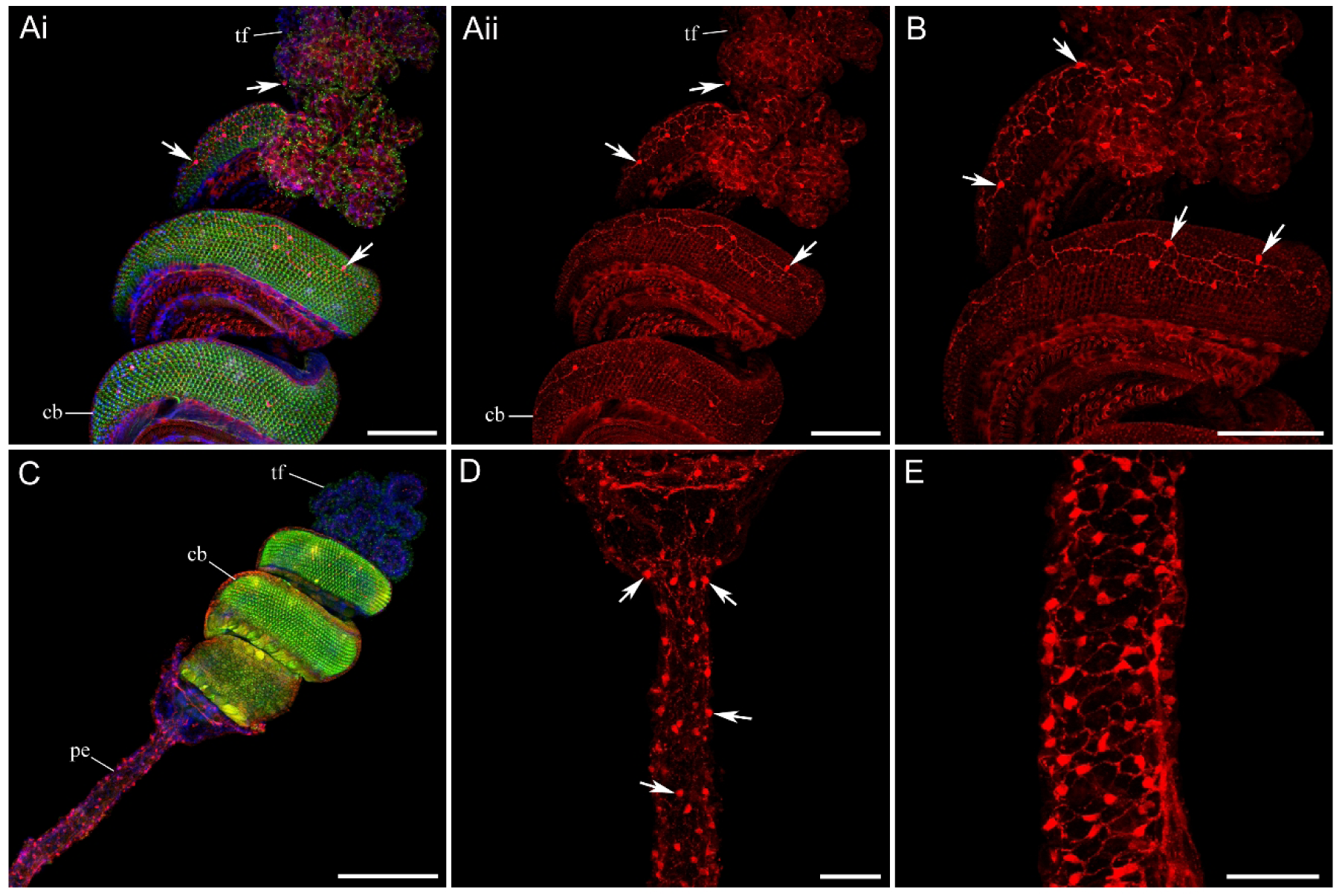
Neural network in the *Nanomia* tentilla. FMRFa IR is red, tubulin IR is green, and DAPI is blue. (Ai) - Cnidoband (cb) and terminal filament (tf) of a single mature tentillum in double-labeling imaging. Note that nematocytes on the outer surface of the spiraled cnidoband are labeled with tubulin AB in green. (Aii) - Red channel with FMRFa IR only. FMRFa-ir neural cell bodies (arrows) are present in both the cnidoband and terminal filament, forming an FMRFa-ir neural network. (B) - Higher magnification shows the FMRFa-ir neural network in cnidoband and terminal filament, with arrows pointing at the neuronal cell bodies. (C) - The entire mature tentillum, including the pedicle (pe), cnidoband (cb), and terminal filament (tf). (D, E) - The pedicle contains a dense FMRFa-ir neural network with numerous immunoreactive cell bodies (arrows). Scale bars: A - 100 µm; B - 100 µm; C - 200 µm; D, E - 50 µm.

Unlike gastrozooids, palpons lack such dense neural system in their main body. Tubulin IR does not reveal any neurons, processes, or networks in the palpon body wall (Fig. 19A-C). FMRFa IR also does not show any scattered neurons or processes in the elongated palpon body (Fig. 19A-C). Only at the base of each palpon, next to its short peduncle connecting to the stem, a plexus or a ring of FMRFa-ir neurons and processes is present (Fig. 19B, C; see Church et al., 2015, Grimmelikhuijzen et al., 1986). Palpon’s single, long and thin tentacle has a chain of FMRFa-ir neurons along its length (Fig. 19D). Nematocysts revealed by tubulin IR labeling are evenly spread throughout the palpacle entire length (Fig. 19D).

**Figure 19.**
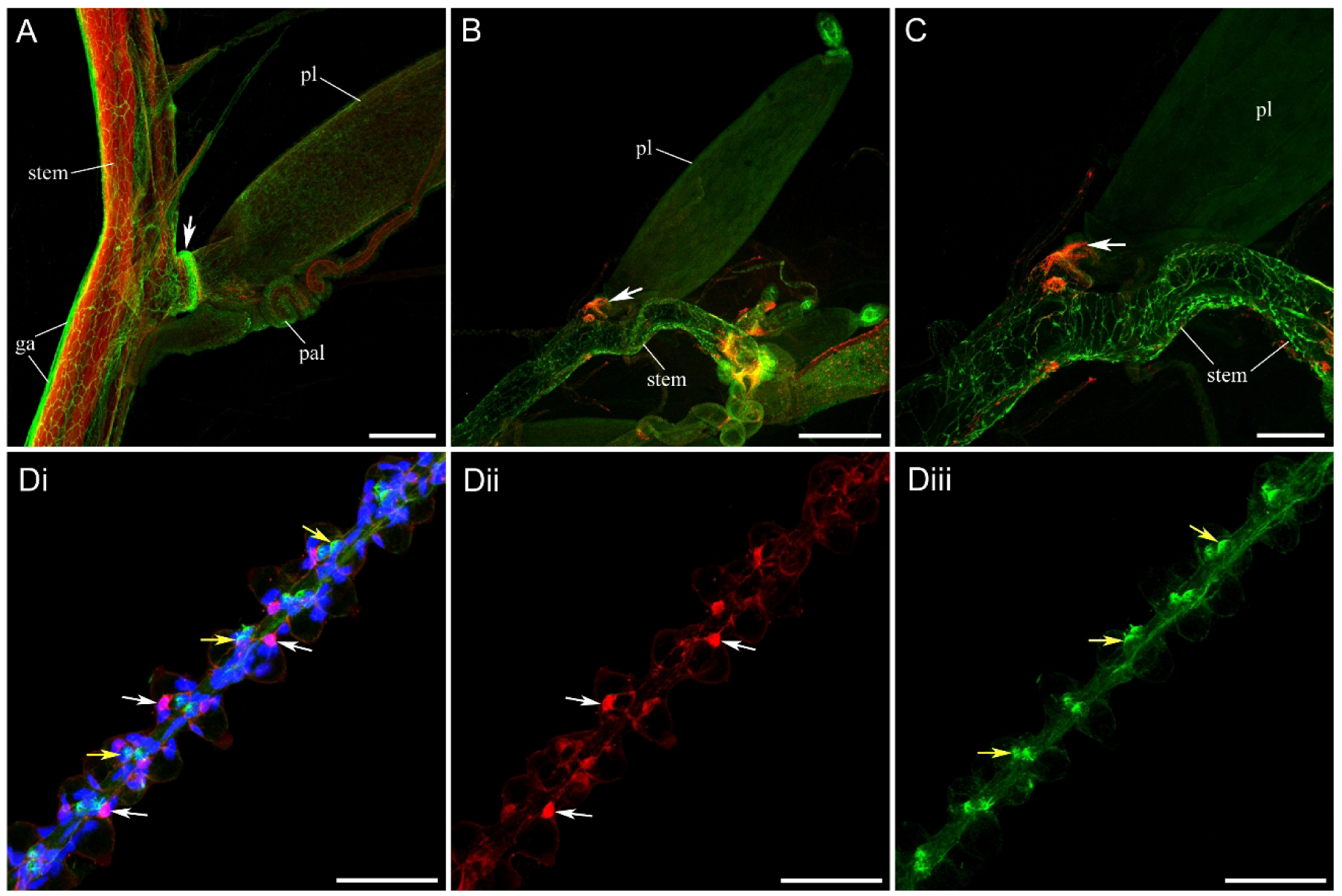
Palpons, or digestive zooids. (A) - Each palpon (pl) attaches to the stem on the ventral side via a short peduncle and has a single palpacle (pal) homologous to the tentacle of a gastrozooid. Tubulin IR (green) labels the stem’s polygonal neural network and a giant axon (ga) on the dorsal side of the stem, while phalloidin labels muscles (red). A narrow ring (arrow) at the base of the palpon, stained by tubulin IR, is similar to the more prominent ring in gastrozooids. (B, C) - Double labeling with tubulin IR (green) and FMRFa IR (red). Neither tubulin IR nor FMRFa IR show any neural elements in the palpon (pl) main body. Only at the palpon base, in the short peduncle that connects palpon to the stem, there is a small FMRFa-ir (red) neural plexus, or a ring, indicated by arrow. (Di) – Long and thin palpacle double-labeled with tubulin IR (green) and FMRFa IR (red). White arrows point at the FMRFa-ir neurons forming a neural chain along its length. Yellow arrows show nematocysts stained with tubulin IR and spread along the entire length of a palpacle. (Dii) – Red channel showing tubulin IR only. (Diii) – Green channel showing tubulin IR only. Scale bars: A, C - 200 µm; B - 500 µm; D - 50 µm.

### Male and Female Gonophores

Male and female reproductive zooids are called gonophores (Fig. 20). A few gonophores connect via a short pedicle to a single stalk and form a reproductive structure called a male or female gonodendron. Each male gonophore has a brightly labeled neural ring at its base, consisting of many small FMRFa-ir neurons and their processes (Fig. 21A, C, D). There are many thin tubulin-ir and FMRFa-ir processes running from the neural ring via a short gonophore pedicle to the stalk and then to the stem, thus connecting each gonophore with the stem nervous system and the rest of the colony (Fig. 21D). In the male gonophores themselves, no FMRFa-ir neurons or processes are noticeable (Fig. 21Dii). Tubulin IR also does not reveal any neuronal structures in male gonophores, although any possible signal from the neural elements could be masked by the brightly labeled with tubulin AB developing sperm, especially at more mature stages (Fig. 21).

**Figure 20.**
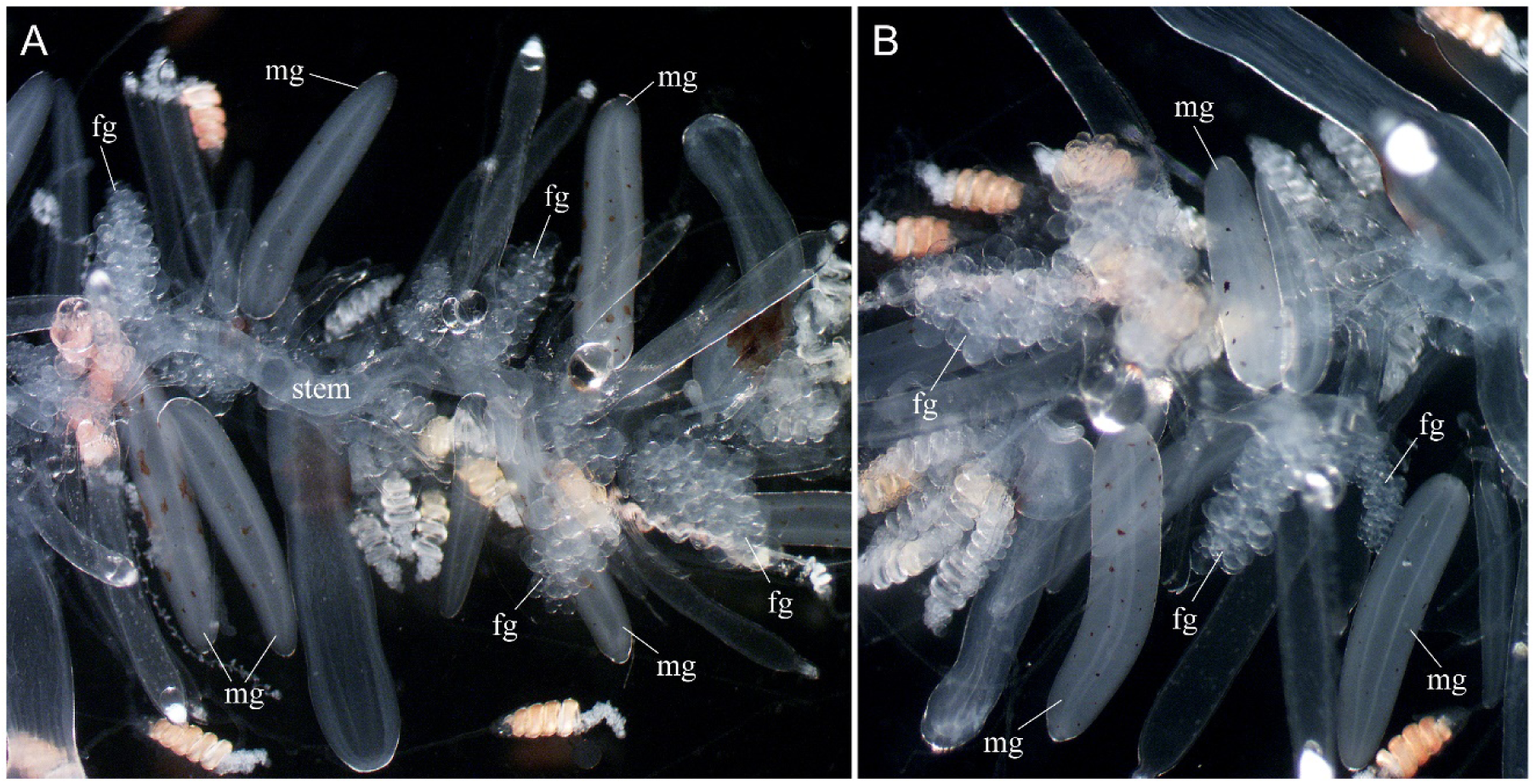
Light microscopy images of the siphosome region, with a focus on the male (mg) and female (fg) gonophores (A, B).

**Figure 21.**
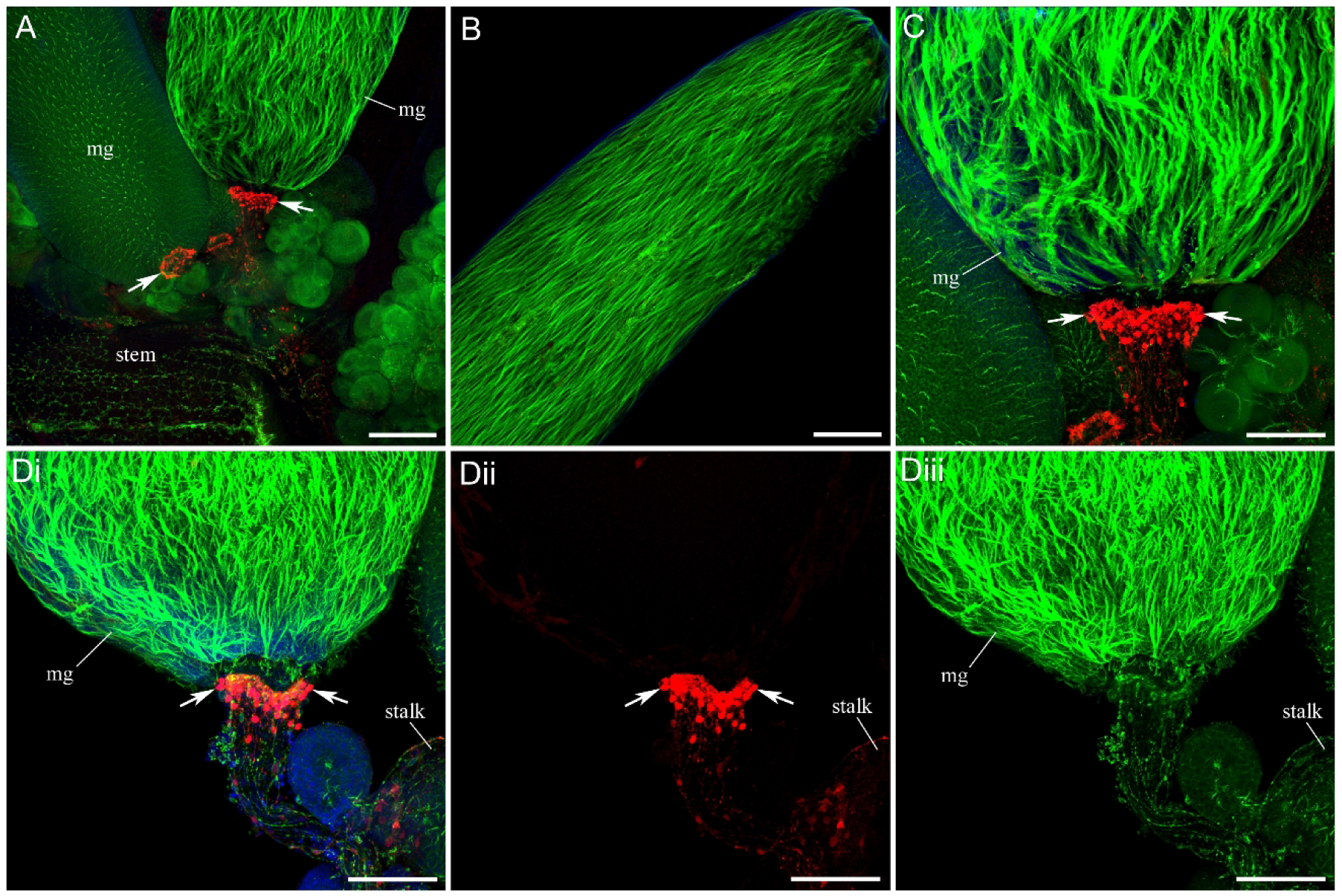
Male gonophores double-labeled with tubulin IR (green) and FMRFa IR (red). (A) - Male gonodendron with at least three male gonophores (mg). Arrows point at the FMRFa-ir neural rings at the base of each male gonophore. Tubulin IR labels numerous mature sperm cells in the gonophore body. They are especially visible in later developmental stages and are not seen in gonophores with immature sperm progenitor cells. (B) - Midsection and distal end of a male gonophore with mature sperm cells. Note the absence of any FMRFa-ir elements. (C) - Higher magnification of the male gonophore (mg) base, with arrows pointing at the FMRFa-ir neural ring. Note numerous FMRFa-ir neural cell bodies in the ring. (Di) - Each male gonophore (mg) resides on a short gonodendron stalk, which then connects to the siphosomal stem. Arrows point to the FMRFa-ir neural ring at the gonophore base. Many thin tubulin-ir and FMRFa-ir processes extend to the stalk and connect the FMRFa-ir gonophore ring to the stem. (Dii) - Red channel with FMRFa IR only. No FMRFa-ir labeling in the body of a male gonophore. (Diii) - Green channel with tubulin IR only. Brightly labeled mature sperm cells in the gonophore can mask any possible tubulin-ir neural elements. Scale bars: A, B - 200 µm; C, D - 100 µm.

As in male gonophores, each female gonophore has an FMRFa-ir neural ring at its base, consisting of many small FMRFa-ir neurons and their processes (Fig. 22B). FMRFa IR also reveals a diffuse neural network of small neuronal cell bodies and their branching processes in the central part of a gonophore, extending throughout its entire length (Fig. 22). Tubulin IR outlines large oocytes located just beneath the outer ectodermal layer.

**Figure 22.**
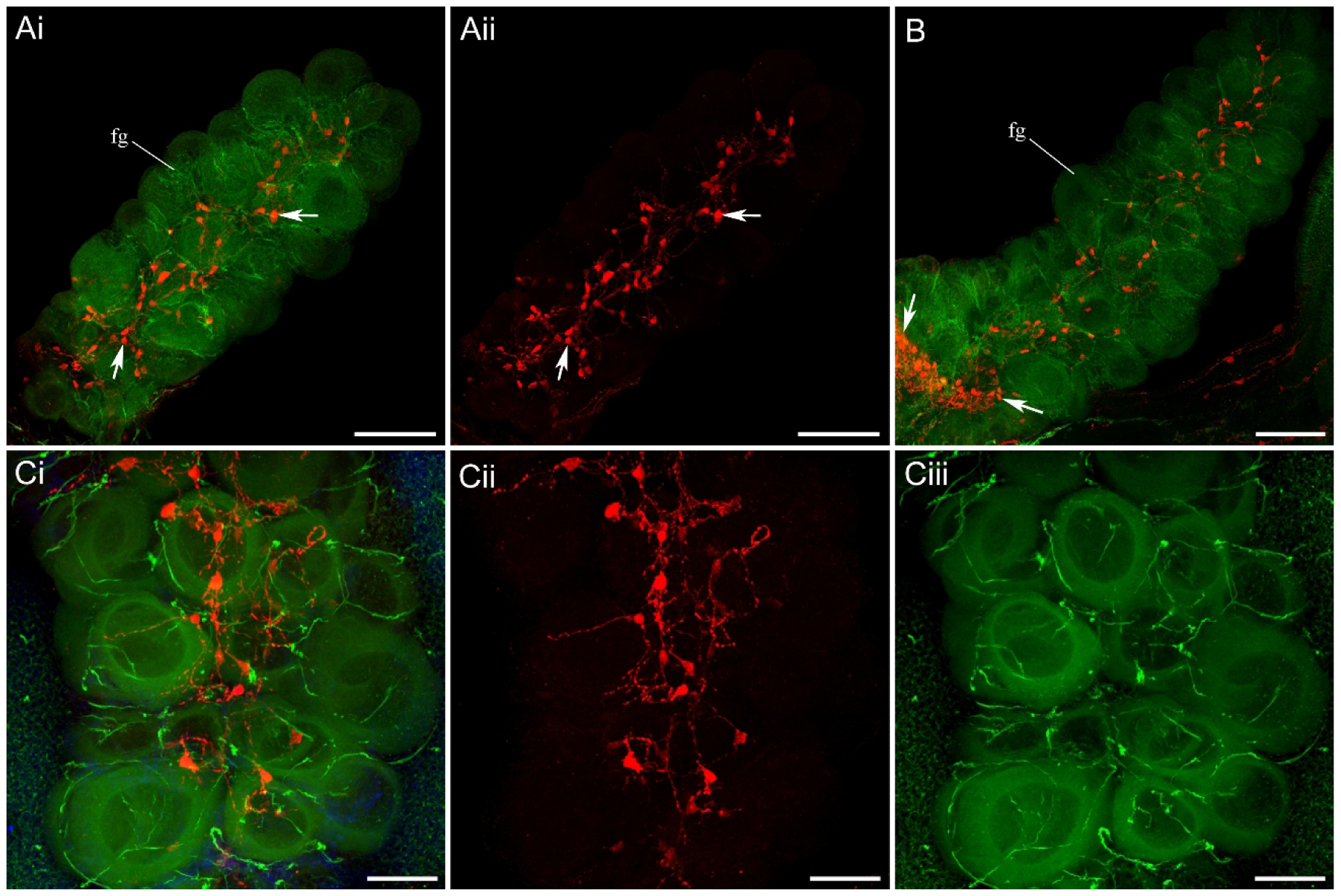
Female gonophores double-labeled with tubulin IR (green) and FMRFa IR (red). (Ai) - Female gonophore (fg). Tubulin IR (green) outlines several oocytes. FMRFa IR (red) labels a diffuse neural network that consists of small neuronal cell bodies and their branching processes. Some immunoreactive neurons are indicated by arrows. (Aii) - Red channel with FMRFa IR only. (B) - Another female gonophore (fg). Note that in addition to a diffuse FMRFa-ir neural network, an FMRFa-ir neural ring (arrows) is present at the base of a gonophore. (Ci) - Higher magnification of a female gonophore at midsection. FMRFa-ir neural cell bodies and their processes are clearly visible in the central part of a gonophore. Tubulin IR outlines several large oocytes and labels many thin filaments, which are presumably sperm cells. (Cii) - Red channel with FMRFa IR only. (Ciii) - Green channel with tubulin IR only. Scale bars: A, B - 100 µm; C - 50 µm.

### Bracts

In *Nanomia*, bracts are specialized, protective zooids spread throughout the siphosome region that resemble small, gelatinous leaves. They primarily serve a defensive function and provide structural support for other zooids. At the base of each bract, a muscular, elongated lamella connects it to the stem, as revealed by phalloidin staining (Fig. 23A-D). These muscle bands (lamellae) in the bracts contain an embedded neural network, labeled by tubulin IR, which is connected to the stem’s polygonal neural network (Fig. 23B-D). In addition to the tubulin-ir network, bract’s lamella contains FMRFa-ir network, which consists of two or more chains of small bipolar FMRFa-ir neurons and connecting processes (Fig. 24A, B; see also Grimmelikhuijzen et al., 1986). No other neural structures are present in the bracts outside of these narrow lamellae and their extensions. The FMRFa-ir and tubulin-ir neural networks in the bract’s lamellae are directly connected to the stem and are continuous with its polygonal neural network (Fig. 24C). The presence of neural networks in the muscular lamellae suggests neural control and a direct connection to the stem polygonal network – an overall integration with the rest of the colony.

**Figure 23.**
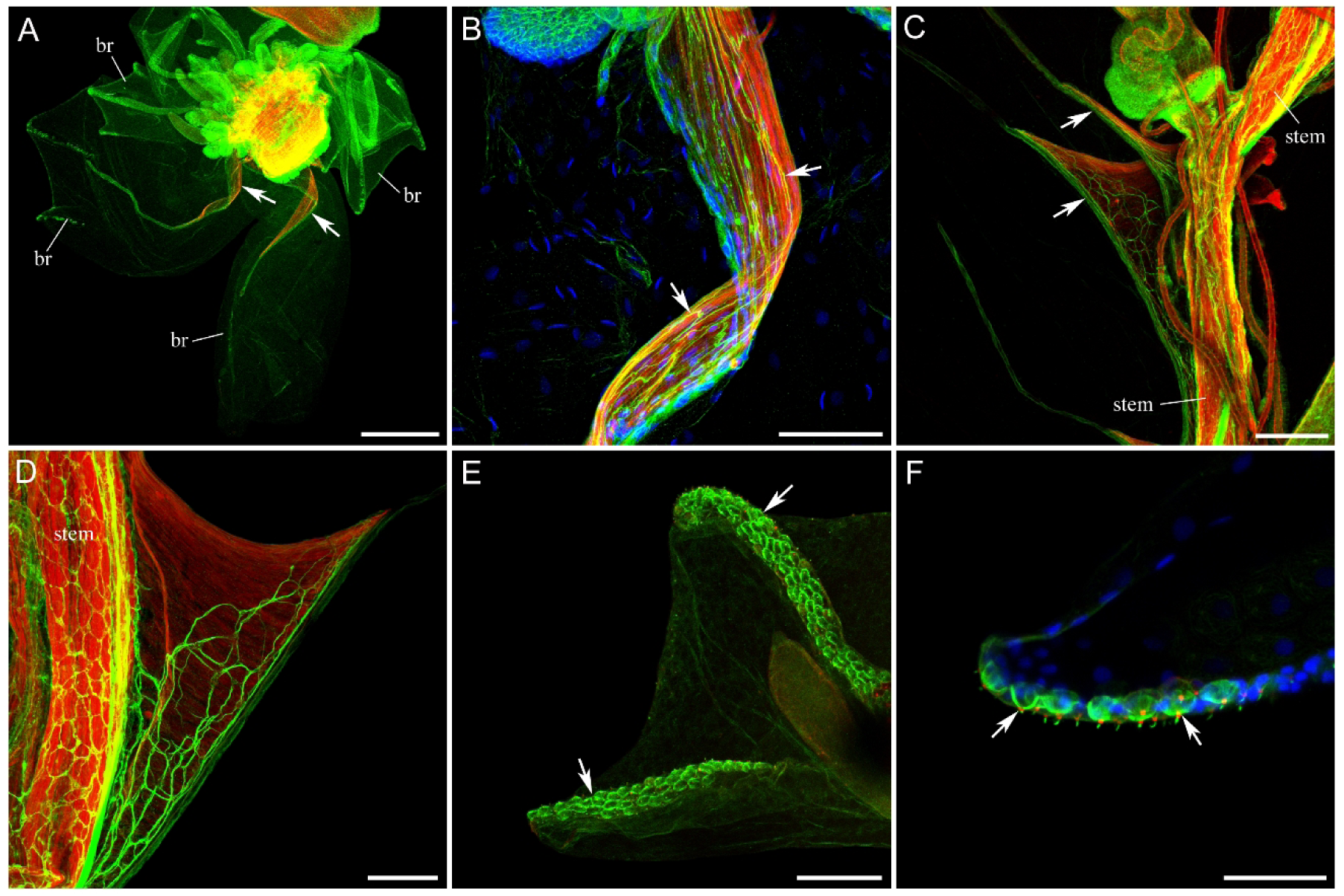
*Nanomia* bracts, gelatinous leaf-like protective zooids, labeled with tubulin AB (green) and phalloidin (red). (A) - Cross-section of the siphosomal region with several small developing and large mature bracts (br). Each large, mature bract has a narrow, muscular lamella (arrows) at its base, labeled with phalloidin in red. (B) - These muscle bands (red) in the bracts contain an embedded neural network (arrows), labeled with tubulin AB (green). (C) - The muscular lamellae (arrows) at the base of the bracts are attached to the stem. (D) - Higher magnification of the bract lamella in longitudinal orientation, with the embedded neural network labeled with tubulin AB (green). (E) - The defensive function of the bracts is provided by three rows of nematocytes (arrows) along the bract ridges. Nematocytes are labeled with tubulin AB in green. (F) - Higher magnification of the bract ridge showing individual nematocytes (arrows), each with a sensory cilium trigger (cnidocil). Scale bars: A - 500 µm; B - 100 µm; C - 200 µm; D, E - 100 µm; F - 50 µm.

**Figure 24.**
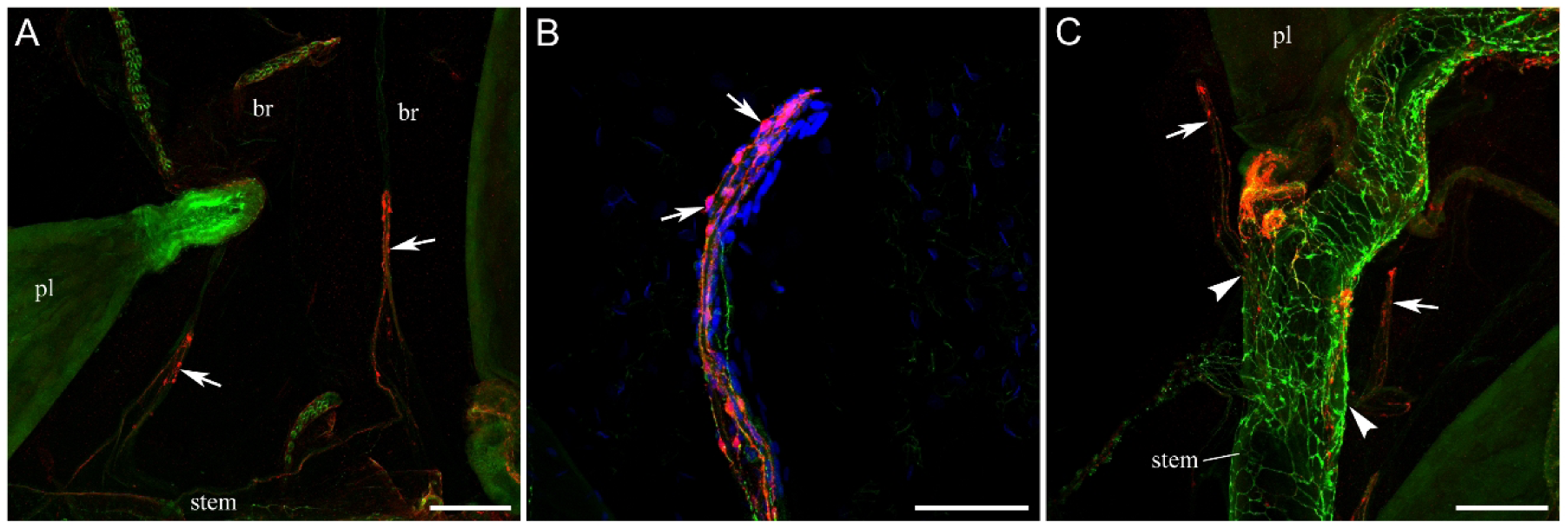
Nervous system in the bracts, double-labeled with tubulin IR (green) and FMRFa IR (red). (A) - The narrow muscular lamella at the bract base (from figure 23A-D) contains not only the tubulin-ir neural network but also FMRFa-ir neural elements (arrows). Note that outside of these lamellae, there are no other neural structures in the bracts. (B) - The neural band at higher magnification: FMRFa-ir neural cell bodies (arrows) and processes, as well as tubulin-ir neural processes (green). (C) - These narrow bands (arrows) at the bract bases, which contain both tubulin-ir and FMRFa-ir neural elements, are directly connected to the stem and its polygonal network (the point of contact is shown by arrowheads). Abbreviations: br - bract; pl - palpon. Scale bars: A, C - 200 µm; B - 100 µm.

The defensive, protective function of the bracts is provided by three rows of nematocytes – two on the side ridges and one along the median line (Fig. 23E). Nematocytes in the bracts and their sensory cilium (cnidocil) are labeled by tubulin IR and are morphologically very similar to the nematocytes on the nectophore ridges (Fig. 23F and 3F).

## DISCUSSION

### General outline

*Nanomia*, as a representative of physonect siphonophores, has one of the most complex colony organizations and behaviors. The long-standing question is “how such composite organisms could mechanistically function effectively and compete on equal terms with unitary zooplankton forms.” (Mackie et al., 1988). Operating as a single individual, unitary organism, siphonophores are **modular** yet more integrated entities compared to conventional benthic cnidarian colonies. In this sense, they are the only ones to have fully exploited and expanded the integrative physiological possibilities of coloniality to their ultimate limits (Mackie et al., 1988). How is this level of integration achieved in siphonophores, and specifically in *Nanomia*? Neural systems in general are essential to the formation of a unitary organism; this is their main functional role. Our mapping of the nervous system in *Nanomia* indicates that all structures are innervated to varying degrees, providing neural continuity across the entire colony, yet with substantial heterogeneity among zooids and between them. The current detailed mapping should take us one step further toward understanding the cellular mechanisms of integration and behavior in the siphonophore colony.

#### Neural system complexity in different zooids reflects the complexity of their function

Individual zooids have their own neural systems that control their function. The complexity of their nervous systems always depends on the complexity of their function and behavior. Gastrozooids that actively capture the prey and initiate digestion have a complex nervous system throughout their entire body, including numerous sensory FMRFa-ir cells in the lip area, polygonal tubulin-ir network and a mash of FMRFa-ir neurons with longitudinal parallel fibers in the hypostome region, and a dense neural network in the tentacles. By contrast, palpons, which function as passive digestion zooids, do not have any neural elements in their thin, elongated bodies. It suggests that their pulsations are endogenous and related to non-neuronal conduction (Mackie, 1965, Spencer, 1971). The gelatinous defensive bracts also do not have any neural elements in most of their body – only in the muscular lamellae at their base that controls their limited movements. However, the pneumatophore (“float” at the colony’s anterior end), which is instrumental for sensory signal processing and locomotion, has very dense neural networks that include tubulin-ir polygonal ectodermal network, numerous FMRFa-ir sensory cells around the pneumatophore pore, and a net of FMRFa-ir neurons and processes throughout its entire body. The nectophores, the medusa-like locomotory zooids that are responsible for all fast escape and slow swimming, also have an extensive neural network representation, including the tubulin-ir nerve ring in control of the swim pattern generation and widely branching sensory-type upper nerve, as well as FMRFa-ir neurons of probably sensory nature around the ring and neural fibers. On the other hand, male gonophores, which serve as a storage for developing sperm cells, lack any noticeable neural systems. Interestingly, there is a network of FMRFa-ir neurons and fibers in the female gonophores. We don’t know their function, but they likely contribute to the active expulsion of individual oocytes upon maturity.

#### Diverse neural elements in the stem serve as a complex coordination center

The stem, which serves as an anchor for all zooids, structurally and functionally integrates the entire colony and has the most complex and dense neural system. The giant axons that run through the nectosomal and siphosomal stem presumably serve to carry signals fast and over long distances – from the anterior pneumatophore or from the distant posterior siphosome region. Giant axons evolved universally and convergently across metazoans, including cnidarians (Mackie, 1973, Norekian and Moroz, 2020a, Roberts and Mackie, 1980) and ctenophores (Mackie et al., 1992, Norekian and Moroz, 2020b) to facilitate the fast propagation of signals over long distances.

The polygonal neural network that covers the entire stem in both nectosome and siphostome regions, is likely capable of carrying a signal over long distances but also operates at local levels, connecting giant axons with the individual zooid attachment areas. The polygonal nerve nets are widely spread in *Nanomia* in different zooids and other hydrozoans, as well as across all ctenophores (Norekian and Moroz, 2020b) indicating a clear convergent trend in their formation. Polygonal architectures might provide cost-efficient, redundant, resilient, and adaptive signal-propagation and integration across uniformed epithelia. In siphonophores, the polygonal network in the stem may represent fused neurons, potentially forming a syncytium not only among themselves, but also with the giant axons, as demonstrated by the injection of Lucifer Yellow (Grimmelikhuijzen et al., 1986). Similar coupling/fusion might be a common feature for various Hydromedusae (Anderson and Mackie, 1977, Spencer, 1981, Spencer and Arkett, 1984).

There are two distinct subpopulations of the polygonal neural network in the stem that match each other in location and polygonal structure. However, one subunit is labeled with tubulin IR only, while the other subunit is double-labeled with tubulin and FMRFa IRs. The presence of a peptidergic transmitter in the second subunit and lack of it in the first might indicate the difference in their functional role, although we do not know exactly what that might be. Differential overlap and partial co-localization of tubulin-ir polygonal networks with peptidergic systems might provide a sufficient foundation for neuroplasticity and modulation of sensory-motor pathways. In all cases, we anticipate a substantial degree of complexity in the co-localization of different transmitters across structures and species, with future directions using various antisera, *in situ* hybridization, or, ideally, *in vivo* imaging.

Additional interesting neural elements in the nectosomal stem are periodic FMRFa-ir neural tracts that consist of small neurons and their processes forming double-chain pathways connecting contralateral cones, serving as anchors for nectophores (Norekian and Meech, 2026). Their functional role is unknown, but it is likely related to the coordination (synchronization or inhibition) of the nectophore activity. These FMRFa-ir double-chain tracts are absent in the siphosomal stem. The characteristic feature of the siphosomal stem is the presence of regular FMRFa-ir transverse rings, which indicate the separation of the siphosomal stem into cormidia - repetitive clusters of specialized zooids (pattern similar to earlier observations by (Grimmelikhuijzen et al., 1986). These FMRFa-ir transvers rings, or collars, can probably either integrate cormidia or isolate them as autonomous units, serving as gatekeepers of information processing in the stem. Interestingly, the FMRFa-ir transvers neural rings (and the corresponding location of longitudinal muscle breaks in the stem, Fig. 15) match the location of the “detachment rings” in the siphonophore species from the suborder Calycophora, in which the cormidia detach as dispersive free-swimming units known as eudoxids during sexual reproduction stage (Manko, Munro and Leclere 2025). *Nanomia* belongs to the suborder Physonecta and does not have a eudoxid sexual reproduction stage and controlled colony fragmentation.

#### Structure of the neural integration between various zooids and the rest of the colony

The most fundamental question remains: how does neural integration between zooids and the central stem occur? We will start from the anterior end of the colony and move through the nectosome region to the posterior siphosome. The anterior float, pneumatophore, appears to be more like an extension of the stem. The polygonal neural network in the stem smoothly transitions into the polygonal network in the pneumatophore without any seam or separation. The tubulin-ir subunit of the polygonal neural network then covers the entire body of the pneumatophore up to the apical pore. Interestingly, the FMRFa-ir polygonal subunit extends only into the base of the pneumatophore and ends there. Thus, these two subunits show a dramatic difference in their architecture in the pneumatophore region. This short FMRFa-ir polygonal network at the pneumatophore base could be mistaken for the stem transverse collar (as suggested by Grimmelikhuijzen et al., 1986). We think this is a FMRFa-ir polygonal network, not a transvers collar typical for the siphosomal stem, which has a different morphological structure.

Nectophore integration with the stem and coordination of nectophore activities is undoubtedly best studied compared to other zooids. The neural connection between the nectophore and the stem occurs via the terminal ganglion at the end of the nectophore lower nerve (Norekian and Meech, 2026). The terminal ganglion is not covered by the ectodermal epithelial layer and directly faces the narrow cleft between the ganglion and the stem cone, which is densely innervated by the stem polygonal neural network with its fibers extending to the cleft from the opposite side. It is not clear how the signal is propagated from the terminal ganglion to the stem and back at the cone junction – whether volume transmission is possible, or if more synaptic-like connections exist. The method used in this study does not allow visualization of individual synapses.

It is important to note another pathway for signal propagation in nectophore coordination and integration. First observations on siphonophores (Mackie, 1960) and later in many other hydrozoans showed that ectodermal (and endodermal) layers consist of electrically coupled epithelial or myoepithelial cells that can propagate electrical signals and participate in multiple levels of coordination, acting in parallel with neural circuits.

There are three types of swimming modes in *Nanomia*: asynchronous swimming during feeding, forward escape swimming triggered by siphosome stimulation, and reverse escape initiated by pneumatophore stimulation (Mackie et al., 1988). Mackie (1964) showed that reverse escape swimming in *Nanomia* depends on signals propagating within the epithelium of the exumbrella ectoderm. It is carried over the entire outer surface of the nectophore by a non-muscular, excitable epithelium (Mackie et al., 1988). While in all nectophores epithelial conductance occurs in the exumbrella, in the stem, it is the endoderm that serves as a conductive medium and not the ectoderm (Mackie et al., 1988). Thus, it was suggested that during reverse escape swimming, the signal travels between the stem polygonal neural network and nectophore ectodermal epithelium via the gap junctions in the cone area (Norekian and Meech, 2026). This epithelial pathway is instrumental in activating Claus’ muscles, responsible for the directional control of jet propulsion in nectophores. At the same time, during slow asynchronous swimming and forward escape swimming, the excitation spreads only between the nectophore terminal ganglion at the end of the lower nerve and the polygonal network in the stem, in neural-to-neural connection (Norekian and Meech, 2026). Thus, the junction site between the stem and nectophores represents a unique decision point stage as a multifunctional, bidirectional gatekeeper that integrates epithelial and neural signals within both condensed and distributed systems.

In the siphosome area, the most noticeable neural element in the location of zooid attachment to the stem is the dense ring of FMRFa-ir neurons and their fibers at the base of each zooid. Our results were comparable to labeling patterns obtained using different antiserum against RFamide sequence (Grimmelikhuijzen et al., 1986) and *in situ* hybridization for *nb*-RFamide (Church et al., 2015). All siphosome zooids (except bracts) are connected to the stem via a short stem branch, which contains numerous neural processes that run from the stem polygonal network into zooids. Above that branch and at the base of each zooid, there is a morphologically distinct ring containing densely packed FMRFa-ir neurons and their fibers. The most prominent neural ring is at the base of gastrozooids, but it is also present at the base of palpons, and male and female gonophores. All neurons in the ring are only FMRFa-ir without any tubulin IR labeling. Some of those neurons could be sensory, based on their morphology – they are bipolar with one cilium-like short projection to the surface, while many others have a simple spherical shape. We do not know the exact role of those FMRFa-ir neural rings, but they could function as a neural relay station between each zooid and the stem, or operate as a gatekeeper of information processing, thus coordinating (synchronizing or inhibiting) the zooid activity with the rest of the colony. Initially suggested hypothesis that the FMRFa-ir neural ring can control the sphincter at the zooid base that closes the continuous flow of the gastric cavity between the stem and the gastric cavities of zooids was rejected, because phalloidin-labeling showed a complete lack of circular muscle fibers that can work as a sphincter. The defensive bracts do not have a neural ring, but the stem polygonal network extends its tubulin-ir and FMRFa-ir neural fibers into the muscular lamella at the base of each bract, which is presumably capable of inducing a range of simple bract movements.

#### The heterogeneity of neural networks

The simultaneous use of two well-known neuronal markers, tubulin AB and FMRFa AB, in our study had several important advantages. The first, and obvious implication was that a much larger population of neural elements was revealed. Many neural networks in *Nanomia* are labeled with only one marker, not the other. For example, the polygonal neural network in the gastrozooid hypostome region is tubulin-ir only, while the mash of sensory neurons in the lip area and scattered small neurons giving rise to longitudinal parallel fibers are FMRFa-ir only. Similarly, in the pneumatophores, the polygonal neural network covering the entire float is tubulin-ir only, while the numerous small cells scattered around its body including the apical pore region are FMRFa-ir only. Such separate labeling of neural populations by tubulin AB or FMRFa AB is not unusual in hydrozoans – for example, we observed similar pattern in jellyfish *Aglantha digitale* (Norekian and Moroz, 2020a). Why many FMRFa-ir neurons are not tubulin-ir is not currently known but presumably reflects the fact that tubulin concentration in those neurons is at a much lower level and does not reach the threshold for tubulin IR identification. Whatever the reason is, using both tubulin and FMRFa IRs reveals about twice as many neurons than one marker alone. In addition to total amount of visualized neurons or networks, the use of two markers allows also to identify and differentiate distinct subpopulations of neural elements in seemingly uniformed single population. For example, as it turned out, the polygonal neural network in the stem consists of two distinct subunits – one is labeled with tubulin IR only, while the other is double-labeled with tubulin and FMRFa IRs. Both subunits have matching morphological structures and architecture and would be totally indistinguishable if not for double-labeling experiments. Another important advantage is identification of the transmitter nature and possible functional role. We used FMRFa AB simply as an additional well-known marker of neural elements in the invertebrate systems. FMRFa AB are not very specific and known to label a wide group of peptides from RFamide family. However, the FMRFa IR indicates that those neurons are peptidergic and could have not only a different transmitter nature, but also a different functional role. For example, peptidergic neural systems are frequently known to have a modulatory role. Although, in *Nanomia* many FMRFa-ir neurons appear to be sensory based on the morphology and location – in the lip area of gastrozooids and around apical pore of pneumatophore, as an example. Nevertheless, neural elements labeled only by tubulin IR can be initially viewed as faster components with more directional transmission – as for example, giant axons, which are labeled by tubulin AB only and are essential for fast escape swimming responses in *Nanomia* (Norekian and Meech, 2026). By contrast, peptidergic neural elements during initial approach could be suggested to have slower transduction and perform more modulatory function.

#### Conclusion and perspectives

We provide evidence of neuroanatomical interactions within all elements of the colony, including contributions of giant axons, stem polygonal nets, and RFamide-ir neural rings at the base of each zooid, as well as describe different subpopulations of neural networks in the body of different zooids. The presented mapping, with more than a dozen morphologically recognized neuronal cell types, facilitates future identification of novel conductive and signaling pathwayFs to decode the cellular basis of behavioral integration within decentralized, broadly distributed networks and non-neuronal elements of these unique superorganisms. However, detailed biophysical and (neuro)chemical mechanisms underlying all interactions (neural and non-neuronal) and integration in siphonophore colonies remain unknown. Despite the fundamental importance of siphonophores as the most advanced and successful colonial organism, the neurobiology of this pelagic group remains largely unexplored. Novel functions of individual zooids are expected to be discovered. For example, nectophores are more than pure locomotory ‘organs’ and include secretory and other cell types. What are the functions of neurons in the unique pneumatophore gas chamber, which is full of toxic carbon monoxide - presumably to control gas release and buoyancy? What is the neural control of reproductive functions, homeostasis, and morphogenesis? What are the behavioral and circuit differences between *Nanomia* and other studied siphonophores? What are the principal differences and similarities between siphonophores and other hydrozoans and cnidarian classes with different body plans? Are there any homologous neurons, circuits, and behaviors?

To move forward, more detailed mapping with *Nanomia*-specific secretory peptides and small transmitters (such as glutamate, GABA, nitric oxide, etc.) is needed, including the possibility of separating synaptic vs. non-synaptic, volume transmission (Moroz, 2021, Moroz, 2023), as well as non-neuronal conductive pathways, as described in other cnidarians (Mackie and Passano, 1968, Satterlie, 2014), with different velocities of conductive pathways (Mackie, 1965, Mackie et al., 1988). The unification of real-time physiological data from natural habitats with single-cell multiomic approaches is highly desirable and achievable, driven by advances in technology and the democratization of novel tools, including mobile floating laboratories (Moroz, 2015), to provide unbiased access to the still-unknown pelagic realm of the world’s oceans.

## Acknowledgments

We thank FHL for their excellent facilities, including the Nikon Laser Scanning confocal microscope. We also thank Dr. Robert Meech for useful discussions of our data. This research was supported by the National Science Foundation grant (IOS-2341882) to LLM.

## Notes

### Competing Interest Statement

The authors have declared no competing interest.

### Summary of Updates

update the orcid.org/0000-0002-7022-1381 for Norekian and the order of authors

